# Dataset structure outweighs method choice in single-cell cell-type annotation

**DOI:** 10.64898/2026.08.28.747622

**Authors:** Oliver Wardhana

## Abstract

Automated cell-type annotation is a prerequisite for nearly all single-cell RNA-sequencing (scRNA-seq) analysis, and the proliferation of annotation tools — spanning marker-based, similarity-based, classical machine-learning, deep-learning, semi-supervised, large language model (LLM), and transformer foundation-model families — has exceeded any existing guidance on how to select among them. Existing benchmarks utilize convenient, well-known datasets in which cell number, class imbalance, cell-type number, and differential-expression strength vary concurrently, so performance cannot be attributed to any dataset property. We benchmarked 63 tools from seven families using a Taguchi L9(3⁴) orthogonal array that varied these four properties independently and extended the comparison under more realistic conditions: real data, cross-platform transfer, public marker databases, LLM annotation, and fine-tuned foundation models. With standardized preprocessing and inputs, the leading cell-level families performed within 0.06 κ of one another, and fine-tuned foundation models averaged only slightly higher. Across families, sequencing platforms, and data types, accuracy was predicted near-linearly by how separable cell types were in a shared expression embedding space, measured as k-nearest-neighbor (kNN) purity (R² = 0.85–0.99). We found that dataset identity accounted for ≈84% of κ variance while tool identity accounted for ≈4%. Strong performers were distributed across families, for example, Seurat label transfer and singleCellNet among reference-based classifiers and mLLMCelltype among LLM annotators. When cell types cleanly separate, many annotation methods suffice; when they do not, algorithmic complexity struggles to compensate for noisy signal. Method selection should therefore prioritize available resources, such as curated references and compute availability, over sophisticated modeling.

## Introduction

Single-cell RNA sequencing (scRNA-seq) resolves cellular heterogeneity at a resolution that reveals rare populations and the composition of tissues [1, 2]. However, a sequenced transcriptomic profile carries no label identity and is not biologically interpretable — each cell or cluster is therefore assigned an identity from its count data. Cell-type annotation is essentially a prerequisite for nearly all downstream analysis, including differential expression, trajectory inference, and cell–cell communication modeling [2, 7]. Annotation has traditionally relied on expert inspection with literature cross-referencing of canonical marker genes across computationally derived clusters. This approach can be subjective, difficult to reproduce, and does not scale: as of 2022, roughly 90% of scRNA-seq studies were still annotated manually [8].

In response, many automated cell-type annotation tools have been developed and span several distinct families [9, 10]. Marker-based tools such as scType [12] and SCINA [13] utilize curated marker sets to score potential labels for cells or clusters. Similarity-based methods, including SingleR [14] and scmap [15], transfer labels from a pre-annotated reference by transcriptional similarity. Supervised classifiers apply classic machine learning methods such as support vector machines, random forests, regularized regression, discriminant analysis, and mixture models [16–20] while deep-learning [21, 22] and semi-supervised [23, 24] models learn nonlinear representations and additionally adapt to the unlabeled query distribution, respectively. Most recently, multi-task transformer foundation models pre-trained on millions of cells [25–29] and generative large language model (LLM) annotators [30–32] have emerged. These families differ substantially in their inputs, statistical assumptions, computational demands, and dependence on external resources, and they seem to increase in complexity over time. Whether that increase delivers better annotations remains unclear.

To date, numerous benchmarking studies have not produced a consistent answer. Support vector machines have been ranked best overall in some comparisons [6, 11]; logistic regression has been shown to match a pre-trained transformer on supervised annotation [33]; and zero-shot evaluations report that foundation models perform inconsistently against simple baselines [34]. The data used in these studies also confound the results: reference–query discrepancy, overlapping populations, finer subtype distinctions, and class imbalance play some role in annotation error [35, 36]. Standard benchmarks use convenient, well-known dataset samples in which cell number, class imbalance, cell-type number, and differential-expression strength co-vary, so the independent contribution of any one dataset property cannot be estimated and performance cannot be causally attributed. Additionally, various upstream steps including clustering, marker derivation, and reference preprocessing may vary between tool pipelines and introduce correlated noise that cannot be separated [38–40].

It is therefore unknown how much of the variation in annotation accuracy is attributable to the annotation method itself and how much is dictated by the structure of the data. To elucidate this, we benchmarked 63 tools from seven families under a Taguchi L9(3⁴) orthogonal array [43, 44], which varies four salient dataset properties (cell number, class imbalance, cell-type number, and differential-expression strength) independently, and we gave each tool the most informative version of the input it utilizes so that performance differences reflect algorithmic capability rather than input quality. We then ran experiments under more realistic conditions — real datasets, transfer between sequencing platforms, public marker databases, LLM annotation, and fine-tuned foundation models — to find which methods perform well and the conditions in which they do (Fig. 1).

**Figure 1.**
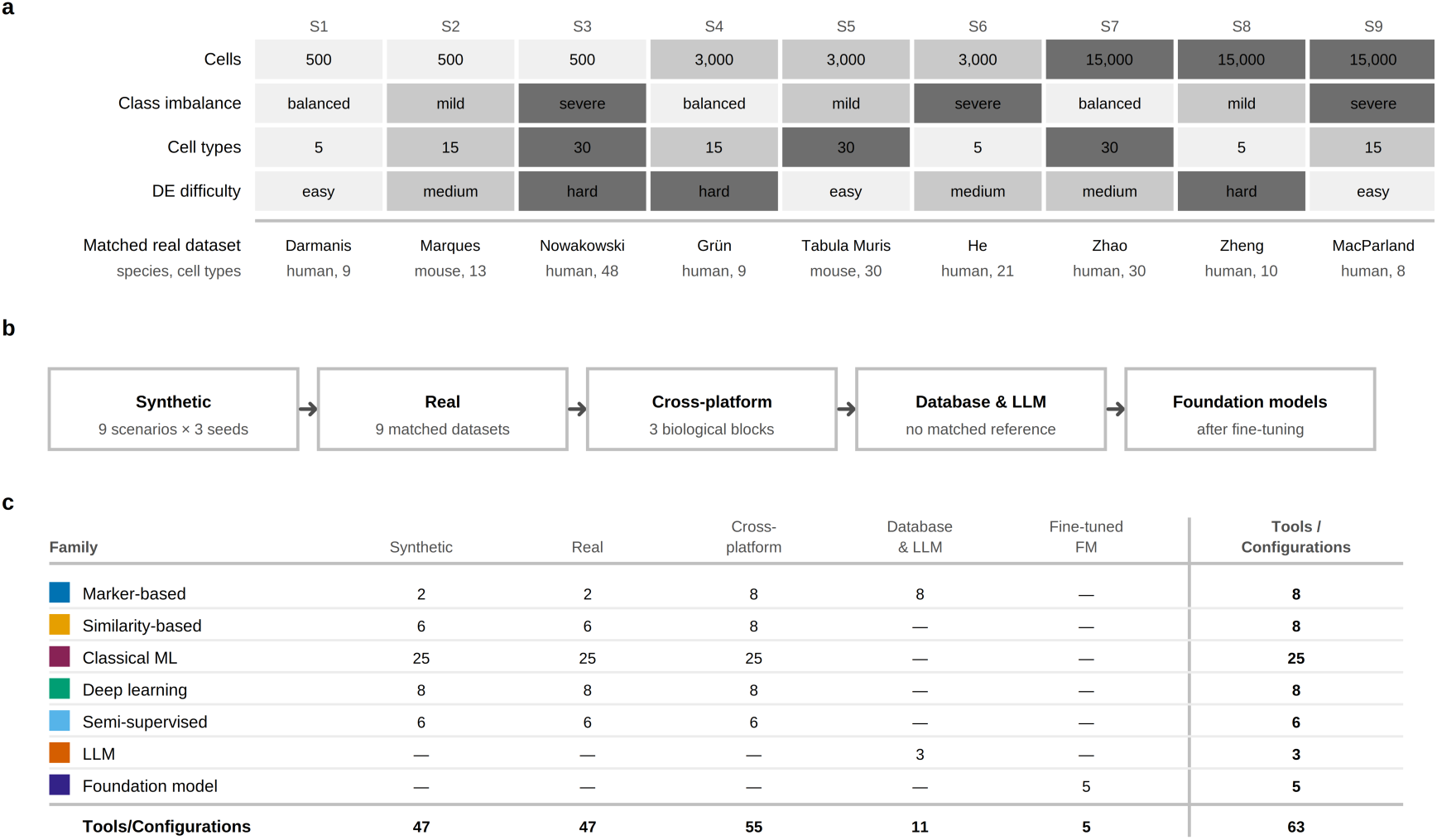
Study design. (a) The nine benchmark scenarios aligned in the Taguchi L9(3⁴) orthogonal design array: four dataset properties (cell number, class imbalance, number of cell types and differential-expression (DE) difficulty) each set to one of three levels across nine synthetic scenarios (S1–S9), shaded by level. The real dataset that matched each synthetic scenario is below (Table 2, Supplementary Table 1). (b) The five evaluation settings, which vary in the degree of experimental control. A standardized Seurat/Scanpy preprocessing pipeline, matched inputs per tool and Cohen’s κ as the metric were held identical across all five. (c) Tool/ configuration counts from each family in each evaluation setting. Cluster-dependent tools are held out of the cell-level comparison (Supplementary Figs 6, 7).

## Results

### Annotation accuracy varies more across datasets than across method families

Family rankings were not consistent between synthetic and real data. On the synthetic data, similarity-based methods led at a mean κ of 0.626, with classical machine learning close behind (0.623), then deep learning (0.604), semi-supervised (0.600), and marker-based (0.558). The four leading families were separated by only 0.026 κ (Supplementary Fig. 1). On real data the order shifted, with similarity-based (0.676) and semi-supervised (0.675) at the top, followed by classical machine learning (0.638), deep learning (0.622), and marker-based (0.484). The four leading families were separated by 0.054 κ (Fig. 2a).

**Figure 2.**
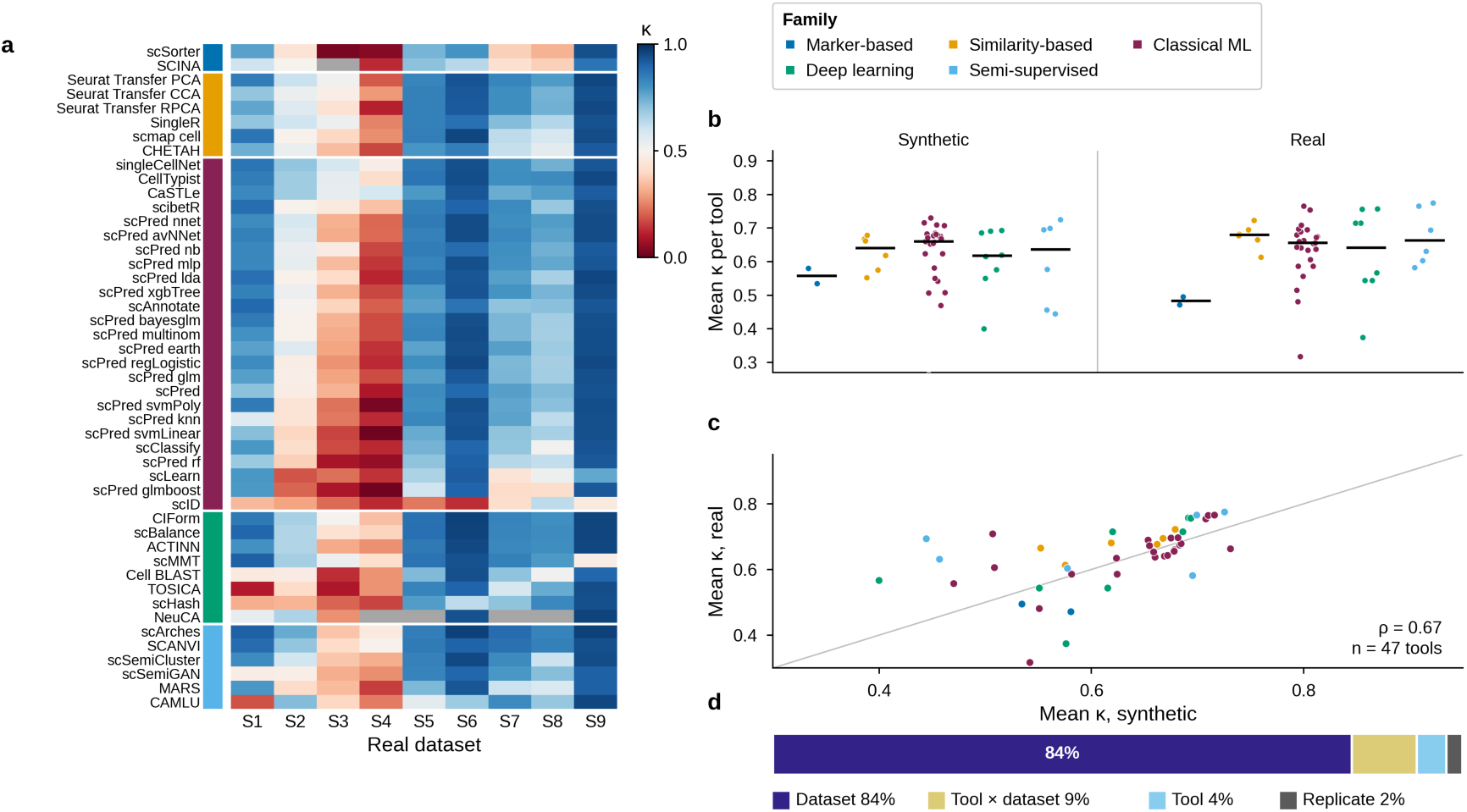
Dataset structure governs annotation accuracy. (a) Per-tool Cohen’s κ across the nine real datasets; 47 tools grouped by family (Supplementary Table 3). Gray cells indicate a failed run with no valid prediction and are considered κ = 0. (b) Mean κ per tool by family, on synthetic and real data; bars are family medians. (c) Mean κ per tool on synthetic versus real data, paired within tool. (d) Variance components from a crossed dataset × tool random-effects model on the synthetic data (n=3). Dataset identity accounts for ≈84% of κ variance, a tool × dataset interaction for ≈9%, tool identity for ≈4% and replicate noise for ≈2%.

Between families, the median of mean κ per tool differed little. Similarity-based tools clustered tightly on both data types, whereas classical machine learning, deep-learning, and semi-supervised configurations were spread more widely. The strongest individual tools were distributed throughout families rather than being concentrated in a particular one. Seurat label transfer (PCA-anchor), singleCellNet, scBalance, and scArches consistently ranked near the top on both synthetic and real data (Fig. 2b). Accuracy seemed to be highly dictated by structural difficulty rather than tool or family choice, as the four structurally ‘easy’ scenarios yielded the highest κ across all families and the two stress-test ‘hard’ scenarios yielded the worst performances overall. Rare cell-type recall showed no consistent family-level pattern on either data type (Supplementary Fig. 3).

Tool-level accuracy on synthetic data scenarios was close to the accuracy on real data scenarios (rank ρ = 0.67; Fig. 2c), with every family gaining slightly on real data except classical machine learning, which was essentially unchanged. Rankings were moderately concordant across the nine scenarios (Kendall’s *W* = 0.42 synthetic, 0.56 real). The reordering could reflect a small interaction between tool and dataset while dataset properties still dominate (Fig. 2d).

### Cell-type separability dominates the variance of κ and predicts annotation accuracy

In a crossed dataset × tool random-effects model fitted to the synthetic data, dataset identity and structure accounted for ≈84% of κ variance, the tool × dataset interaction for ≈9%, tool identity for ≈4%, and replicate noise for ≈2% (Fig. 2d). A decomposition over the four continuous dataset descriptors, pooled across tools and including a tool-identity term, also agreed: kNN purity carried the largest unique share on both synthetic and real data (η² = 0.71 and 0.39) and tool identity a small share (0.06 and 0.14) (Supplementary Fig. 5a,b). When this model was fitted on each family separately, purity’s attribution ranged from 0.63 to 0.83 on synthetic data and 0.19 to 0.57 on real data, with the other dataset properties contributing η² ≤ 0.05 in every family (Supplementary Fig. 5c,d).

This dominance seems to arise through a tight and linear relationship when plotting family-mean κ against kNN purity at the dataset level in every family on both data types — R² = 0.88–0.98 on synthetic data and 0.85–0.99 on real data, with Spearman ρ between 0.92 and 1.00 (all *p* ≤ 0.001; Fig. 3a; Supplementary Fig. 1 for the synthetic data). This near-linearity is strong and consistent in the cell-level annotation performance data and possibly motivates the use of kNN purity as a pre-annotation diagnostic.

**Figure 3.**
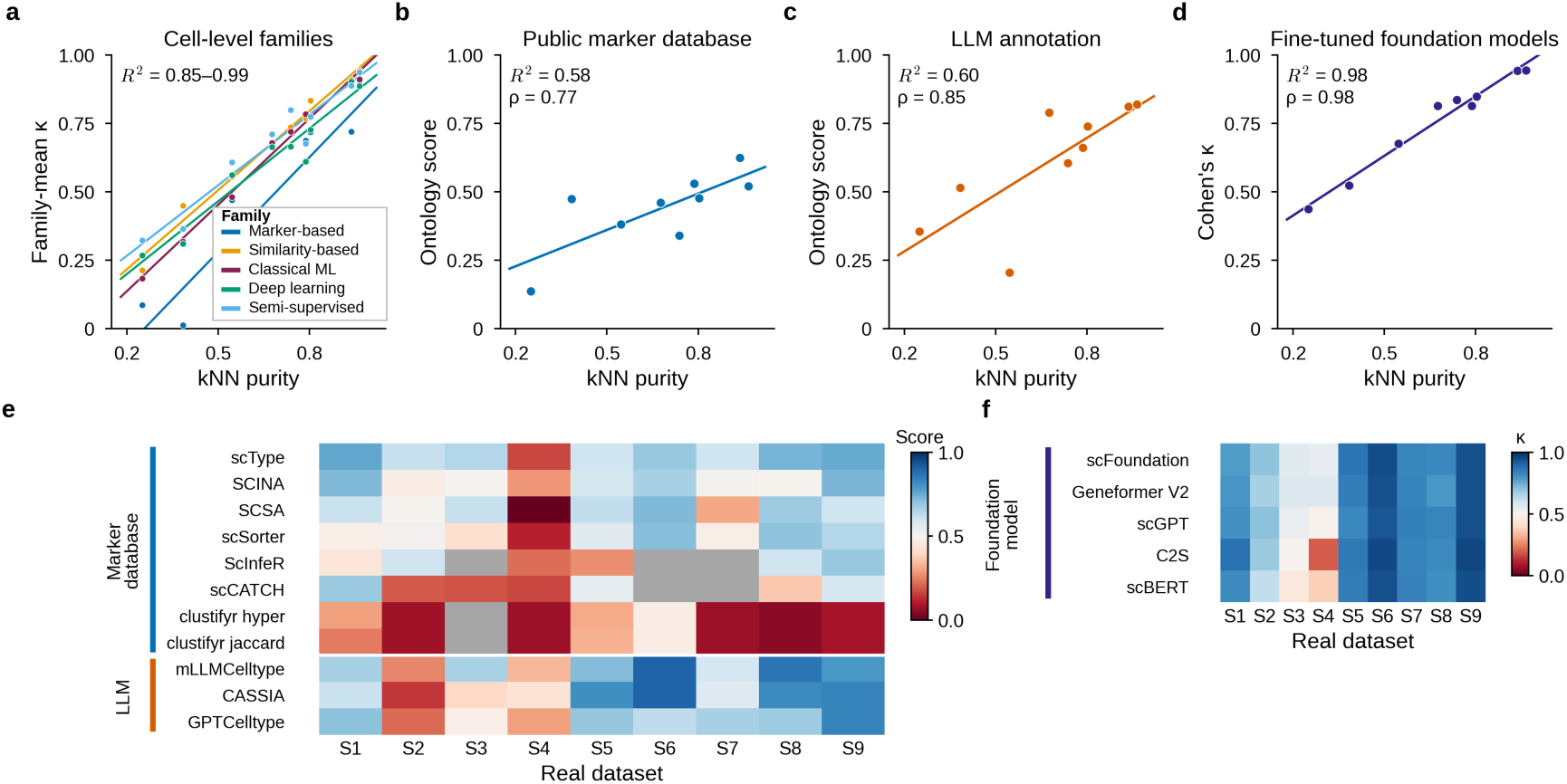
The separability–accuracy linear relationship holds in every data and experimental regime. (a) Family-mean Cohen’s κ versus kNN purity across the nine real datasets for the five cell-level families; each family is fitted separately (n = 9 datasets per fit). (b) Mean ontology-matched score versus kNN purity for the eight database-driven marker tools; scores are the mean of three independent scorers on a 0/0.5/1 scale. (c) The same for the three LLM annotators. (d) Mean Cohen’s κ versus kNN purity for the five fine-tuned foundation models. (e) Per-tool scores across the nine real datasets for the database-driven and LLM families; gray cells indicate failed annotation runs. (f) Per-model κ across the nine datasets for the five foundation models. Fits in (a)–(d) are ordinary least squares at the dataset level; R² = 0.85–0.99 across the five families in (a), and 0.58, 0.60 and 0.98 in (b), (c) and (d), with Spearman ρ = 0.77, 0.85 and 0.98 respectively.

Out of the four dataset properties, purity (a proxy for differential expression) is the dominant driver of κ. Differential-expression difficulty was the dominant driver of κ variance among the four design factors (categorical η² = 0.667; continuous proxy 0.706), ahead of cell-type number (0.096), cell count (0.074), and class imbalance (0.004). Fitting the same decomposition with purity as the response variable reproduced that ordering — differential-expression difficulty accounted for η² ≈ 0.722 of purity variance, cell-type number ≈ 0.146, cell count ≈ 0.130, and imbalance ≈ 0.002 (Supplementary Fig. 4b,c). Purity is a consistent, measurable surrogate for differential-expression difficulty that describes embedding separability and correlates with annotation accuracy.

Synthetic and real data agree in the direction and slope of these effects but differ in absolute magnitude. Synthetic datasets formed smooth, well-separated islands whose separability tracks the differential-expression parameter directly, whereas real datasets show irregular, noisy topologies (Supplementary Fig. 2). Absolute κ is therefore inflated on the less noisy synthetic data in the ‘easy’ scenarios (S1, S5, S6, S9).

### Cross-platform annotation transfer is still governed by biological difficulty rather than tool

Testing annotation performance when the reference and query come from different sequencing platforms showed that accuracy tracked the difficulty of the datasets rather than the platform contrast or the tool used. Pooled across the 55 tool configurations, median κ was 0.99 for CellBench (n = 220 tool × trial values; IQR 0.92–1.00), 0.90 for the pancreas block (n = 110; IQR 0.82–0.93), and 0.62 for PBMCbench (n = 165; IQR 0.52–0.70). The performance difference between blocks exceeded any within-block platform contrast even though detected-gene overlap was comparable across blocks (Jaccard 0.53–0.75). Profiling placed PBMCbench as a difficult dataset, with lower median library size, fewer detected genes per cell, and lower kNN purity. Within a block, no systematic differences in performance distinguished droplet-to-droplet, droplet-to-nanowell, and droplet-to-plate transfers (Supplementary Fig. 8).

### Cell-type separability dictates performance even without exact reference data

Replacing the derived marker list with a single public database (CellMarker 2.0 [71], tissue-subsetted by scenario) lowered marker-based accuracy as expected. Among the eight database-driven tools, scType scored highest (mean ontology-matched score 0.618; Fig. 3e) with a family mean of 0.438 which is below the 0.498 mean κ that the same marker tools reached with derived markers (Supplementary Fig. 7). Although the two metrics are not directly comparable, the direction of the performance change is as expected when substituting a marker list derived from the original dataset with an imperfectly matched external knowledge source.

LLM annotators outperformed the database-driven marker tools on all nine datasets with a family mean of 0.611. Within the family, the multi-agent and consensus configurations had better performance compared to single-call: mLLMCelltype 0.643, CASSIA 0.613, GPTCelltype 0.578. LLM tool failures emerged on datasets whose ground-truth labels do not map cleanly onto Cell Ontology [72] terms such as the Marques dataset (S2). This is a limitation of free-text scoring rather than of supervised label transfer.

The linear purity-to-performance relationship existed clearly in both regimes — Spearman ρ = 0.77 (*p* = 0.016, R² = 0.58) for the database-driven family and ρ = 0.85 (*p* = 0.004, R² = 0.60) for the LLM family (Fig. 3b,c). Similar to earlier trials, the other three dataset descriptors showed weaker associations. The reduction in R² compared to trials with derived inputs is perhaps consistent with the noise that external knowledge sources and free-text annotation introduce.

### Fine-tuned foundation models are the strongest family and follow the same purity– accuracy relationship

Fine-tuned foundation models’ performance matched but did not broadly exceed that of the strongest conventional tools. The five models spanned a narrow performance band — scFoundation 0.777, Geneformer V2 0.772, scGPT 0.767, C2S 0.741, scBERT 0.739 — for a family mean of 0.759 which is above every cell-level family mean on real datasets (0.484– 0.676). The best individual conventional tools reached a κ of 0.85–0.92 on the structurally ‘easy’ datasets which is above the foundation-model family mean on those same datasets (Fig. 3f).

Considering the architectural and pre-training diversity of the five foundation models, the narrow performance spread suggests that at the scales tested, architecture matters less for accuracy than the data regime and the effect of fine-tuning. The cell-type separability-to-performance relationship was tightest in this family as mean κ across the five models was near-perfectly linear with kNN purity (Spearman ρ = 0.98, *p* < 10⁻³; R² = 0.98; Fig. 3d).

### Computational cost relates to workflow accessibility rather than accuracy

Accuracy showed no meaningful association with either runtime or peak memory at the tool-by-dataset level (|ρ| ≤ 0.14; Supplementary Fig. 9). However, runtime and memory overhead differed across families by orders of magnitude: marker-based and similarity-based methods completed in seconds to minutes, classical machine learning in minutes, and deep-learning and semi-supervised methods in tens of minutes to hours, with peak memory following the same trend (Supplementary Figs 10 and 11). Cost varied as much within a regime as between regimes: median runtime per dataset spanned 0.3 s (clustifyr Jaccard) to 1,855 s (scSorter) across the eight database-driven marker tools, and within the three LLM annotators a single-call prompt ran in a median of 5 s against 844 s and 1,763 s for the multi-model consensus and multi-agent pipelines (Supplementary Fig. 12). Cell count governed runtime to an extent for similarity-based and classical machine learning methods (Supplementary Fig. 13). Within the LLM family, runtime scaled with the number of cell types (ρ = 0.85, *p* = 0.004, R² = 0.72) which is reflective of additional inference calls per additional cluster. Foundation models’ runtimes scaled near-linearly with cell count (log–log slope ≈ 0.96, R² = 1.00), spanning seconds to hours depending on model and dataset.

In practice, families are better distinguished by accessibility and integrability rather than their performance capabilities (Fig. 4). Where a labeled reference and GPU capacity are available, fine-tuned foundation models gave the highest mean agreement (κ ≈ 0.76). Where a reference exists but compute is limited, similarity-based methods are competitive (κ ≈ 0.68). Where no reference is available, LLM annotators labeled existing clusters with comparable agreement (score ≈ 0.61).

**Figure 4.**
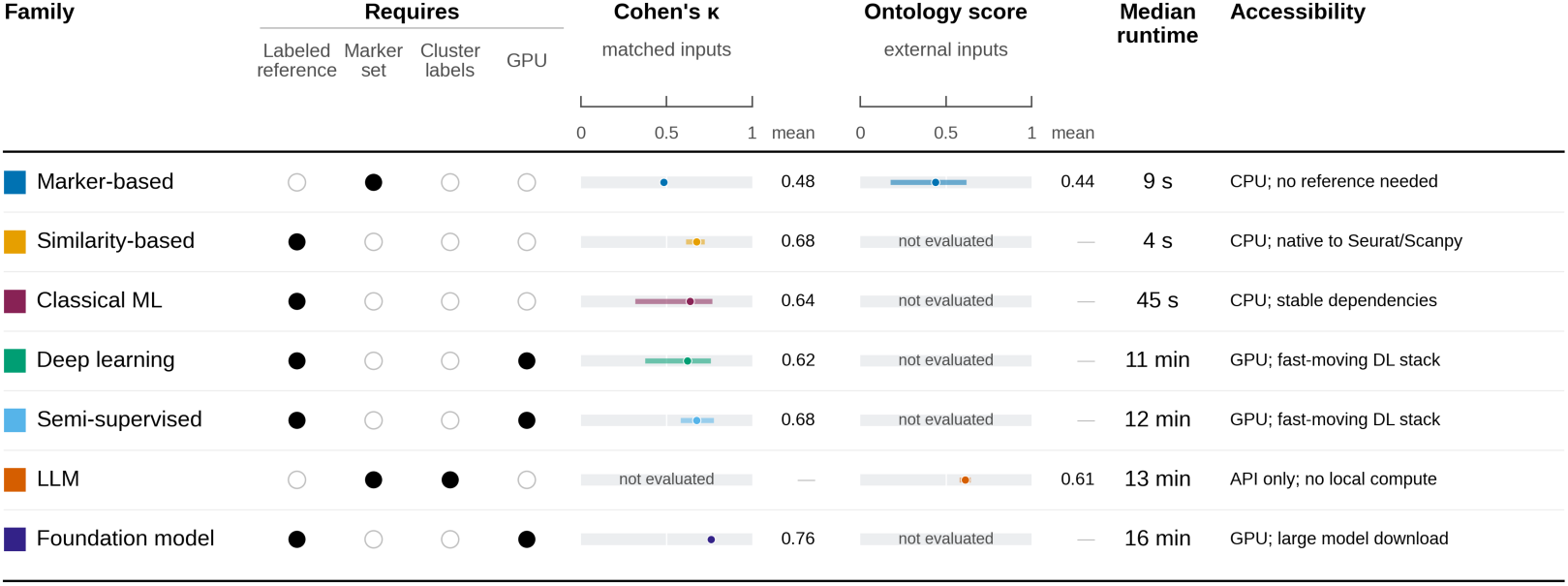
Method selection is constrained by available resources rather than annotation power. (Fig. 1c, Supplementary Table 3). κ is the metric for the real data, cell-level trials with a matched reference or derived markers, the ontology score is the metric for real data, cluster-level trials utilizing a public database or producing free text annotations. For Cohen’s κ and ontology score, the line spans the range and the mean is indicated. Median runtime is the per-(tool, dataset) median on real data, end-to-end for the tool call.

## Discussion

Across various experimental settings, differences between the cell-level tool families were small compared to the differences between datasets: the four families represented by more than two tools performed within 0.06 κ of one another on real data, while per-dataset means across all tools spanned 0.21 to 0.91. Fine-tuned foundation models averaged slightly higher than any cell-level family, though that margin is small beside the dataset-to-dataset spread. Accuracy tracked how cleanly cell types separate in expression space, as kNN purity accounted for the largest share of the variance in κ in every family and on both synthetic and real data. This relationship was close to linear throughout, even being observed in cross-sequencing platform annotation, in the withdrawal of an exactly matched reference, in the substitution of an external marker database, in free-text annotation, and in fine-tuned foundation model annotation. This contrasts with earlier benchmarks that identified particular algorithms or architectures that showed broadly superior performance [6, 11] and supports other studies that called attention to the effect of dataset properties [35, 36]. We attribute this discrepancy to confounding between dataset properties. When dataset properties co-vary, algorithmic differences may look pronounced, but when those properties are held orthogonal by study design, the differences seem to shrink and the variance is attributed to the data instead.

Given these results, it may be more appropriate to ask how well any cell-level method can be expected to perform given a dataset’s structure rather than asking what tool is best. When cell types separate cleanly, many tools reach high accuracy and the choice can be made on software accessibility, hardware requirements, and how well a tool fits an existing pipeline. Where populations overlap heavily, no tool performed well, and manual interventions such as re-embedding, batch integration, or expert curation may be necessary. kNN purity can be used as a near-linear pre-annotation diagnostic by utilizing the labeled reference set or a clustered query set, subject to the caveats in Supplementary Note 3.

On accessibility, similarity-based methods and manual curation dominate published workflows largely because they are well maintained, integrate with scRNA pipelines easily, and run quickly on ordinary CPU hardware. On the other hand, deep-learning, semi-supervised, and foundation-model tools depend on GPU access and on volatile deep-learning libraries, with several tools requiring extensive environment pinning or failing to install cleanly due to dependency conflicts. Considering the putative accuracy ceiling set by the data, accessible tools are more practical for bioinformaticians. Pre-trained, tissue-specific classifiers such as CellTypist [18] additionally extend accessibility through the distribution of pre-trained models, though with a fixed label vocabulary.

The language-model tools are among the newest and more experimental of the families, and their results suggest a shift in what annotation is. LLM annotators need neither a curated reference atlas nor hardware requirements beyond an API. However, they consume cluster-level input and thus inherit upstream clustering errors, they recall rare and tissue-specific types less reliably, and their free-text output maps imperfectly onto controlled cell-type vocabularies. Still, they represent a paradigm shift in which cell-type annotation is moving from closed-vocabulary classification toward semantic reasoning over markers, literature, and ontologies.

Foundation models are also more experimental and they achieved the highest family mean on real data. While corpus-scale pre-training seemed to raise the performance attainable, most of the benefit we observed came from fine-tuning rather than pre-training scale. This is consistent with reports that untuned foundation models do not reliably beat simple baselines [34]. More testing is needed to understand scaling laws in the setting of scRNA-seq and answer several limitations attached to this study (Supplementary Note 2).

### Conclusions

The choice among existing cell-level annotation methods appears to matter less for accuracy than the structure of the data being annotated. Therefore, tool selection can be reasonably made on practical grounds based on the availability of a reference, the available compute, and integration with an existing pipeline. Our results suggest that a cell-type separability check can be run before analysis to determine the expected performance of automated annotation tools should they be used. Lastly, for method development, accessibility, active maintenance, and interoperable label vocabularies are near-term gains that are as consequential as advances in classifier design and are perhaps more immediately useful for high-throughput labs. Standardizing annotations against the Cell Ontology [72] would make annotations easier to compare across studies and would also improve how free-text methods are evaluated. The results here do not argue against new algorithms but suggest that annotation accuracy and accessibility depend mostly on the data and on the environment and infrastructure around these tools.

## Methods

### Study design

We evaluated 63 annotation tools under five experimental settings which vary in their degree of deployment realism: a factorial simulation using an orthogonal design, a real-data panel matched to the synthetic scenarios, annotation transfer between sequencing platforms, annotation driven by a public marker database and by LLMs, and fine-tuned foundation models. A standardized preprocessing pipeline, common input rules, and benchmarking harness were held constant.

### Tool selection and inclusion criteria

Distinct configurations of the same tool were counted separately (scPred’s 17 kernel variants, scmap’s two prediction modes, clustifyr’s two similarity modes); the eight database-driven marker tools are the same physical tools used in the marker-based comparisons and were not double-counted. Three rules governed which of the screened tools were included. Tools requiring closed proprietary infrastructure were excluded. Cluster-dependent tools (those that require an upstream clustering step) were excluded from the cell-level family comparison. Tools that produced degenerate or unstable output were omitted. All 63 included tools, their family, mechanism, inputs, and inclusion status are listed in Supplementary Table 3.

### Preprocessing

All datasets were passed through a standardized pipeline implemented in R (Seurat package) [52, 53] and mirrored in Python (Scanpy package) [70]. Cells expressing fewer than 200 genes were removed and genes detected in fewer than three cells were discarded. Count data were log-normalized and the top-2,000 highly variable genes were selected by Seurat’s variance-stabilizing transform [52]. The matrix was subset to those features, scaled to unit variance with values clipped at 10, and reduced by principal component analysis retaining 50 components. The first 30 PCs were used for any neighborhood-based analyses. Gene identifiers were aligned to a common human symbol convention before this pipeline (Supplementary Method 1).

### Standardized inputs

Each tool was given the most informative form of the input(s) it consumes, each derived from the original dataset. Three input classes are defined.

The *reference matrix* is a labeled training partition obtained by stratified 80/20 split on the original dataset. This expression matrix shares the gene space, cell-type vocabulary, and cell-type proportions of the test partition.

The *marker list* is the top 20 differentially expressed genes per cell type, computed with FindAllMarkers() (Wilcoxon rank-sum, only.pos = FALSE, min.cells.group = 3), filtered to positive markers and ranked by avg_log₂FC. No p-value filter was applied.

The *cluster-identity vector* is the ground-truth cell-type label vector, supplied to tools that require cluster identity as an input. We therefore assumed that a ground-truth cell type derived from the training-set partition matched a cluster.

For the cross-platform transfer experiments, the reference set was drawn from a dataset independent of the test partition source. In database-driven annotation, the derived marker list was replaced by tissue-specific subsets of a public marker database (CellMarker 2.0). Because these inputs are more informative than what is typically available (i.e., label set, marker list, clustering, gene coverage, batch structure, and class frequencies do not completely match the query set), the performance figures reported in this study are upper bounds. Performance differences between tools are mostly unconfounded and reflect algorithmic capability rather than input quality.

### Benchmarking harness

All tools were run through a single framework, which received standardized training and test objects and returned per-cell predictions and ground-truth labels. R-based tools operated on Seurat objects and Python-based tools operated on AnnData objects. Runtime and peak system memory were recorded around each tool’s core annotation call which includes any model training and inference. Memory was captured via peakRAM in R tools and parsed from native system reporting for Python tools.

### Factorial simulation under an orthogonal design

Synthetic data were generated with Splatter [41] via splatSimulate(), with nGenes = 10000, lib.loc = 8.5, lib.scale = 0.5, and dropout.type = “none” [42] chosen so that the simulated libraries fall in the range of a 10x Chromium PBMC dataset [3]. Every scenario was profiled before use: sparsity was 85.6–86.8% and median library size 4,746–4,920 UMIs per cell across the nine scenarios, both within the 10x range. Differential-expression difficulty was set by de.prob(0.15 / 0.08 / 0.04) and de.facLoc (1.50 / 0.80 / 0.30) for easy / medium / hard, giving measured mean up-regulation fold-changes of 4.8×, 2.4×, and 1.6×. This corresponds to clearly distinct populations, moderately differentiated subtypes, and subtly differentiated cell states respectively.

The Taguchi L9(3⁴) orthogonal array [43, 44] that was used varied four dataset properties independently across nine scenarios: cell count (500 / 3,000 / 15,000), class imbalance (balanced; mild, largest-to-smallest ratio R = 5; severe, R = 20), cell-type number (5 / 15 / 30), and differential-expression difficulty as above (Table 1). This orthogonal construction permits unbiased estimation of each property’s effect on performance. Three seed replicate simulations were generated per synthetic dataset scenario. Batch, donor, doublet, and cell-cycle variation are unmodeled in this design. Note that the orthogonal array is a main-effects design, so there is a standard assumption of negligible interactions across the values tested.

**Table 1.** Taguchi L9(3⁴) orthogonal array. Each row defines one scenario, instantiated with three replicates (seeds 42, 123, 999). *R* is the largest-to-smallest cell-type count ratio.

| Scenario | Cells | Imbalance | Cell types | DE difficulty |
| --- | --- | --- | --- | --- |
| S1 | 500 | Balanced | 5 | Easy |
| S2 | 500 | Mild ($R = 5$) | 15 | Medium |
| S3 | 500 | Severe ($R = 20$) | 30 | Hard |
| S4 | 3,000 | Balanced | 15 | Hard |
| S5 | 3,000 | Mild ($R = 5$) | 30 | Easy |
| S6 | 3,000 | Severe ($R = 20$) | 5 | Medium |
| S7 | 15,000 | Balanced | 30 | Medium |
| S8 | 15,000 | Mild ($R = 5$) | 5 | Hard |
| S9 | 15,000 | Severe ($R = 20$) | 15 | Easy |

### Real-data validation panel

Forty-four candidate datasets drawn from the scRNAseq package and other widely used cell-type-resolved datasets were screened [3, 74–77]. Nine were selected by consensus profiling alignment to match the nine synthetic dataset scenarios. The candidates were profiled on: total cell count, cell-type count, minimum cluster size, normalized Shannon entropy of the cell-type distribution, dropout percentage, median UMIs per cell, a biological-coefficient-of-variation dispersion proxy, PCA silhouette width, kNN purity, and mean differential-expression strength (π) [78]. Each was assigned to its nearest scenario under three schemes: the full ten-metric profile, projection onto the four design metrics, and PCA fitted on the real-data profiles. The assignments were combined by Borda-count rank aggregation, with the highest aggregate score taken as the structural match. Assignments reflect the best match across the full profile despite some agreement on individual axes (Table 2, Supplementary Tables 1, 2).

**Table 2.** Real-data validation panel and Borda-consensus mapping to the synthetic scenarios. Profiling values describe the dataset object actually benchmarked (the variant selected by Borda consensus, downsampled where applicable) and are the values used for variance decompositions and structure–performance fits.

| ID | Dataset | Species / tissue | Cells | Cell types | Min. cluster | Norm. Shannon entropy | kNN purity |
| --- | --- | --- | --- | --- | --- | --- | --- |
| S1 | Darmanis Brain 2015 [79] | Human cortex | 466 | 9 | 16 | 0.88 | 0.79 |
| S2 | Marques Brain 2016 [80] | Mouse oligodendrocyte | 2,993 | 13 | 44 | 0.93 | 0.55 |
| S3 | Nowakowski Cortex 2017 [46] | Human developing cortex | 2,978 | 48 | 9 | 0.93 | 0.38 |
| S4 | Grün Pancreas 2016 [81] | Human pancreatic islets | 1,728 | 9 | 96 | 0.93 | 0.25 |
| S5 | Tabula Muris FACS 3k subset 2018 [74] | Mouse multi-tissue (FACS) | 2,984 | 30 | 42 | 0.94 | 0.80 |
| S6 | He Skin 2020 [77] | Human skin | 2,990 | 21 | 1 | 0.53 | 0.94 |
| S7 | Zhao Immune Fine subset 2020 [50] | Human immune subtypes | 14,986 | 30 | 55 | 0.89 | 0.74 |
| S8 | ZhengSort 20 subset 2017 [3] | Human PBMC (FACS) | 20,000 | 10 | 2,000 | 1.00 | 0.68 |
| S9 | MacParland Liver Broad 2018 [45] | Human liver | 8,444 | 8 | 20 | 0.71 | 0.96 |

### Cross-platform transfer

Cross-platform transfer and batch effects were evaluated using matched datasets that share cell-type composition across different sequencing platforms (Supplementary Table 2). CellBench [75] covered 10x v2, CEL-seq2, and Drop-seq on identical lung cancer cell-line mixtures. The pancreas block paired Baron [83] (inDrop) with Segerstolpe [84] (Smart-seq2). PBMCbench [76] matched a 10x v2 reference against Drop-seq, inDrops, and Seq-Well on matched PBMC samples. For each, tools were trained on one platform and tested on another.

### Database-driven marker and LLM annotation

For the database-driven trials, the marker list was replaced by tissue-specific subsets of CellMarker 2.0 [71] (Supplementary Method 2). Performance of the marker and cluster-dependent tools using derived markers rather than the public database is reported as a baseline (Supplementary Figs 6 and 7). LLM tools received the top 20 differentially expressed genes per cluster (marker list) as the primary input. Both families were evaluated without a train–test split.

Because both families emit free text rather than closed-vocabulary predictions, agreement was quantified with a 0 / 0.5 / 1 ontology-matched score (0 = no match, 0.5 = partial, sub-, or supertype match, 1 = match) averaged across three independent LLM scorers (Gemini 3 Flash, Claude Haiku 4.5, GPT-5.4-nano).

### Fine-tuned foundation models

Five foundation models (scBERT, scGPT, scFoundation, Geneformer V2, C2S) were fully fine-tuned and tested on the nine real datasets using the identical stratified 80/20 partitions. Gene vocabularies are model-specific and genes absent from a model’s token vocabulary were dropped prior to tokenization. Evaluation was restricted to real data under fine-tuning, as these models were pre-trained on real transcriptomes and it is known that untuned models perform poorly against simple baselines [34].

### Metrics and statistical analysis

Annotation accuracy was quantified with Cohen’s κ [85], which corrects for chance agreement and is appropriate for multiclass evaluation under class imbalance; κ = 0 is the baseline [11, 35, 36]. Secondary metrics also recorded included: macro F1, the multiclass Matthews correlation coefficient (Gorodkin formulation [86]; Supplementary Note 1), runtime in seconds, and peak resident-set memory in megabytes.

Cell-type separability was measured as kNN purity — it is calculated for each cell as the fraction of its k = 15 nearest neighbors in the 30-component principal-component embedding that share its ground-truth label, averaged over all cells. Purity is a downstream proxy for separability rather than an independent cause.

The contribution of dataset properties to κ was characterized by a Type III (marginal-after-all-others) η² decomposition over four continuous dataset descriptors — log₁₀ cell count, cell-type number, Shannon entropy of the cell-type distribution, and kNN purity. Type III sums of squares give each predictor its unique contribution while the others are held fixed, thus the attribution of variance is order-independent. We found the descriptors were close to orthogonal in this panel (maximum pairwise r = 0.36; VIF 1.15–1.19) (Supplementary Fig. 4a; Supplementary Method 4). The decomposition was first computed in a pooled manner across all tools, with a tool-identity term included and then calculated separately within each tool family, where no tool term is fitted.

We additionally fitted a crossed two-way random-effects model (κ ∼ scenario × tool) on the synthetic data to separate tool-by-dataset interaction from replicate noise. Variance components were estimated by the method of moments. To validate the orthogonal design and establish the pathway by which the L9 design factors act, the same categorical decomposition was applied with kNN purity as the response rather than κ (Supplementary Fig. 4b).

### Computing environment

R pipelines (Seurat marker and reference methods, statistical analysis, figure generation) ran under R 4.5.1 with Seurat 5.0 and Bioconductor 3.21 on an Apple M1 Pro laptop. Non-foundation-model Python tools ran on a shared workstation with a 56-core Intel Xeon E5-2690 v4 (2.60 GHz), 500 GB RAM, and an NVIDIA GeForce RTX 5060 GPU. Foundation-model fine-tuning ran on Google Colab: scBERT, scGPT, scFoundation, and Geneformer V2 on NVIDIA L4 (22.5 GB VRAM), and C2S on NVIDIA RTX PRO 6000 Blackwell Server Edition (102 GB VRAM); C2S differs from the other four in backbone and starting checkpoint (Supplementary Method 3).

## Author Contributions

OW: Conceptualization, investigation, methodology, software, data curation, formal analysis, validation, visualization, and writing – original draft, writing – review & editing.

## Competing Interests

The author declare no competing interests.

## Funding

Glynn Honors Program from University of Notre Dame (OW).

## Data Availability

All real datasets analyzed were previously published, public, and obtained through the Bioconductor scRNAseq package or from data distributed with their original publications (Supplementary Table 1). Reference code used in this study is available at https://github.com/ owardhana/scRNA-Annotation-Benchmarking-Runbook.

## Notes

### Competing Interest Statement

The authors have declared no competing interest.

### Summary of Updates

Major updates throughout entire text and all figures - organization, rewriting, new figures for streamlining and clarification purposes. Author list updated for confidential reasons.

