## Supplemental Information and Materials for "Dataset structure outweighs method choice in single-cell cell-type annotation"

### Supplementary Figures

Figures reporting eta-squared give the proportion of total variance attributable to a factor. Residual fraction is shown explicitly. Unless stated otherwise the modeling unit is the per-(tool, dataset) mean of the response.

*Supplementary Figure 1 — Annotation accuracy on the synthetic data*

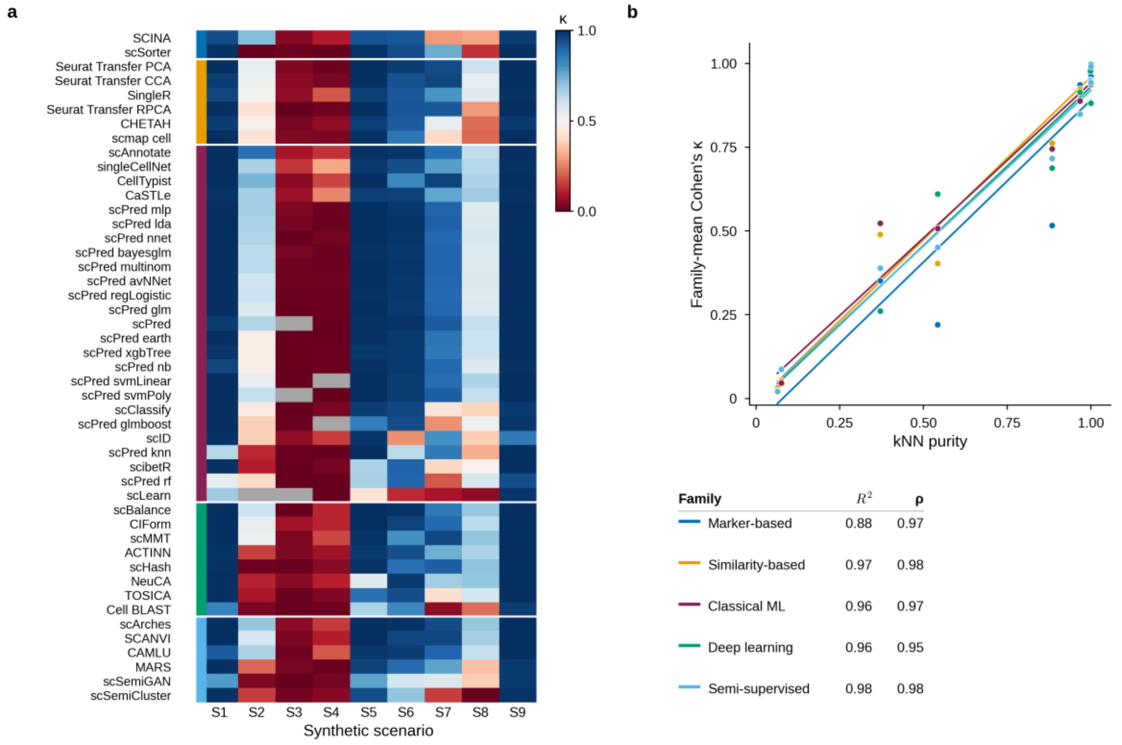


**(a)** Per-tool Cohen's κ across the nine synthetic scenarios, the synthetic counterpart of Figure 2a. Each cell is the mean κ over three replicate seeds (42, 123, 999). Gray cells are failed runs and are treated as κ = 0. **(b)** Family-mean κ against dataset kNN purity across the same nine scenarios ($n$ = 9), the synthetic counterpart of Figure 3a.

The same trend in performance seen on the real data is seen here: the two hard, high-cell-type scenarios (S3, S4) are poor for almost every tool regardless of family, while the four structurally easy scenarios (S1, S5, S6, S9) are strong for almost every tool. Within any one column the spread across the 47 tools is small next to the spread across columns within any one row.

The purity relationship is linear in every family, with $R^2$ = 0.88–0.98 (semi-supervised 0.979, similarity-based 0.966, classical machine learning 0.963, deep learning 0.961, marker-based 0.884) and Spearman ρ between 0.95 and 0.98. Fits are slightly tighter than on the real data scenarios ($R^2$ = 0.85–0.99, Fig. 3a) and span a wider kNN purity range. This result supports the purity–performance trend seen on the real data.

*Supplementary Figure 2 — Paired synthetic and real UMAP embeddings, all nine scenarios*

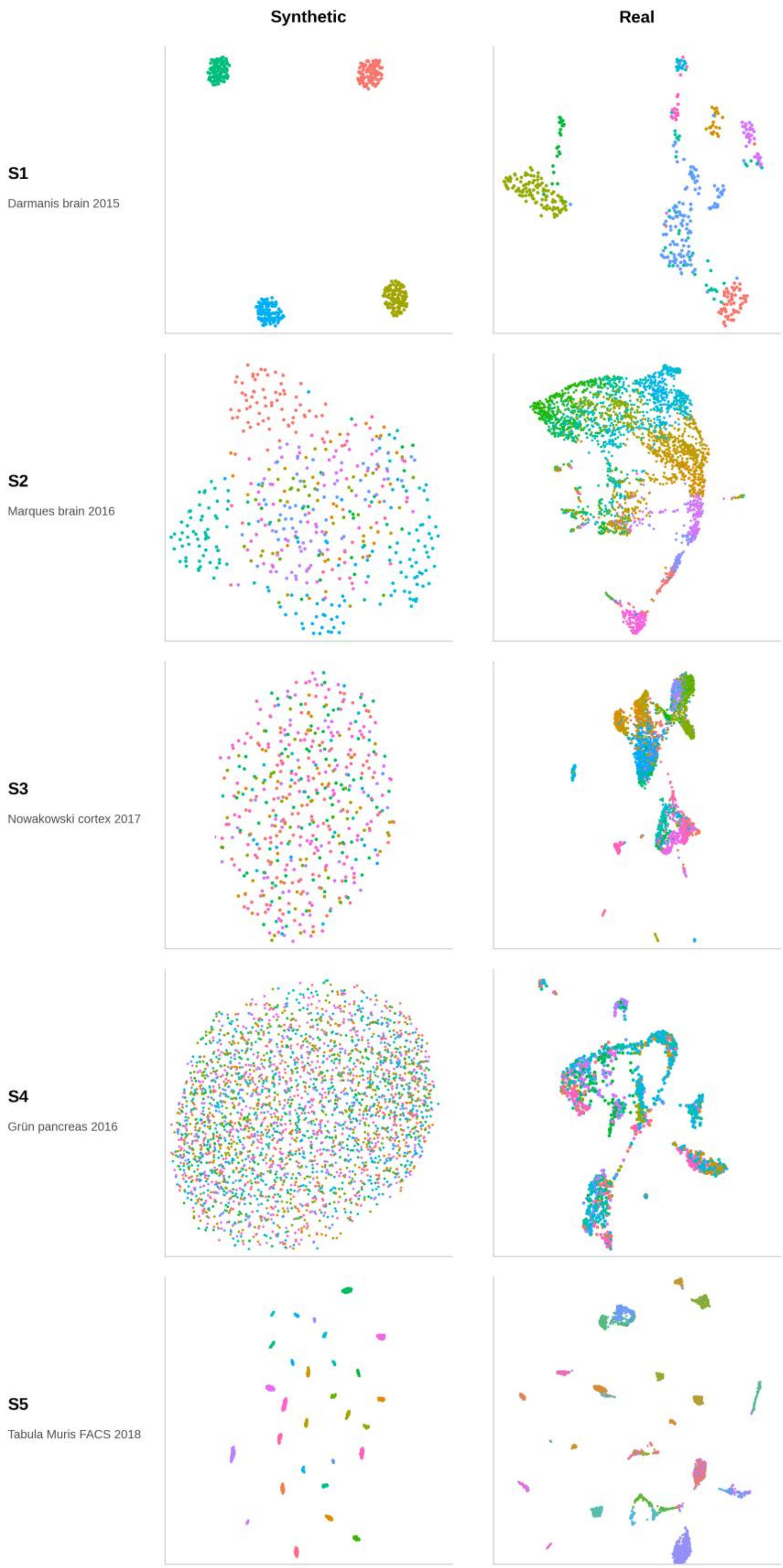

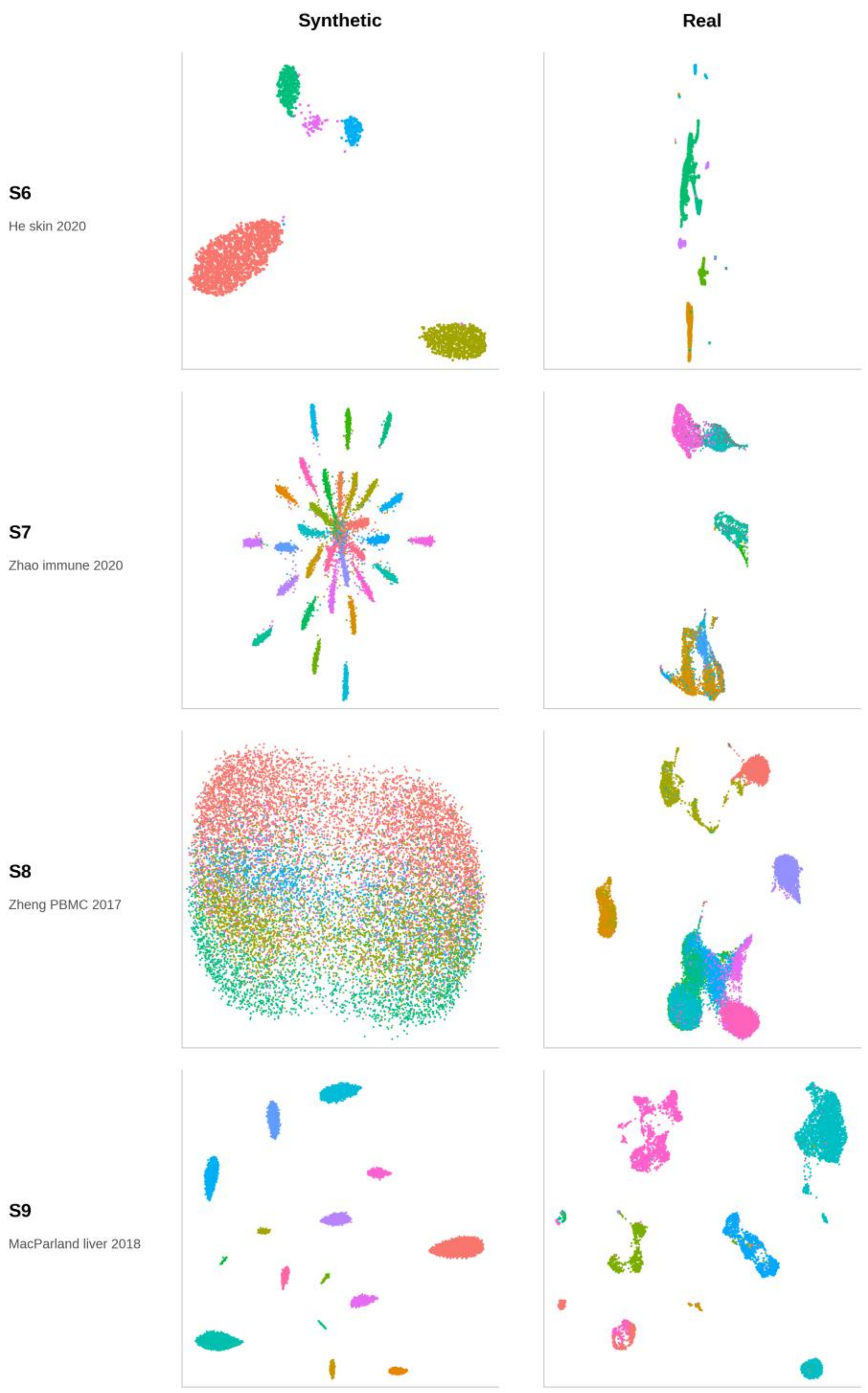


UMAP embeddings of each L9 synthetic scenario (left column) paired with its matched real dataset (right column), colored by ground-truth cell type. The datasets have between 5 and 48 cell types.

The synthetic embeddings show the characteristic Splatter geometry — smooth, oval-like islands whose separability tracks the DE-difficulty setting: clean islands in scenarios 1, 5, 6, 7 and 9; Venn-style overlap in 2 and 8; near-complete overlap in 3 and 4. Real data embeddings show the irregular characteristics of noisy biological data. The geometric idealization of the synthetic data explains some κ inflation on synthetic data. In both data regimes, it is evident how DE-difficulty level and embedding separability are captured quantitatively by kNN purity.

#### *Supplementary Figure 3 — Per-family rare cell-type F1*

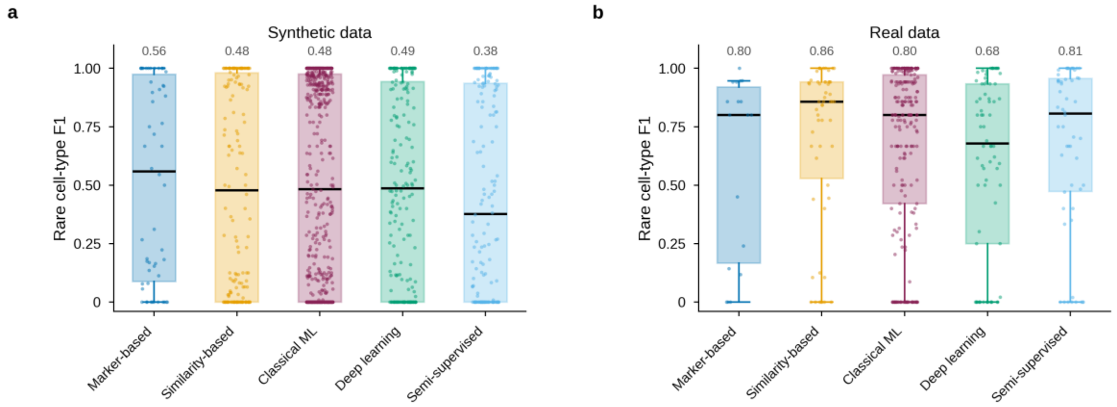


Rare cell-type F1 by family across the synthetic and real dataset scenarios. Rare cell types are defined per dataset as those constituting less than 5% of total cells. No consistent family-level pattern emerges even between the real and synthetic data scenarios. This implies that rare-cell identification is governed by the separability of the rare population rather than the complexity of the tool.

*Supplementary Figure 4 — Orthogonality of the continuous-proxy framework*

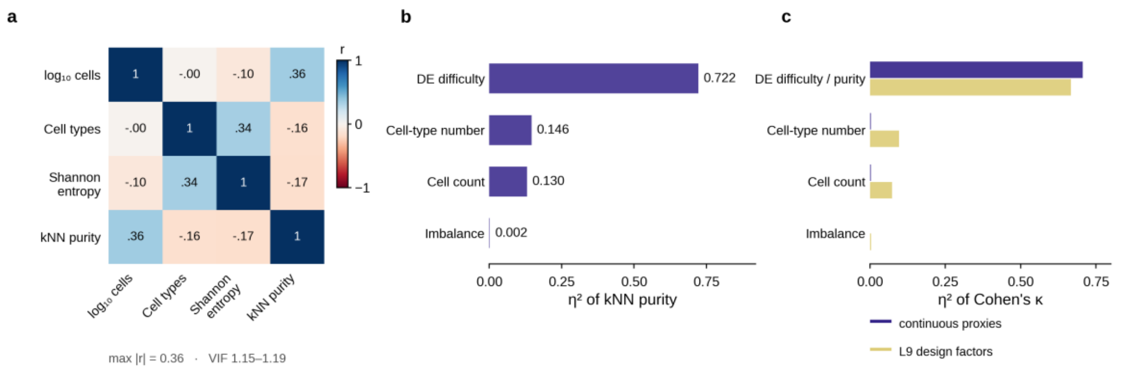


Diagnostic for substituting continuous profiling proxies for the discrete L9($3^4$) variables on the synthetic data (see Supplementary Method 4). All decompositions are Type III.

- **(a) Multicollinearity screen.** Pearson correlation heatmap between the four continuous proxies across the nine synthetic scenarios. Maximum off-diagonal $r = 0.36$ ($\log_{10}$ total cells vs kNN purity); VIFs bounded 1.15–1.19, well below the severe-collinearity threshold of 5.0.
- **(b) kNN purity decomposition.** Variance decomposition of kNN purity across the four design factors at their categorical levels. DE difficulty alone accounts for $\eta^2 = 0.722$ of purity variance, 0.146 for cell-type number, 0.130 for cell count and 0.002 for imbalance.
- **(c) Continuous proxy validation.** Synthetic data variance decomposition on the four L9 factors at their categorical levels and as continuous proxies. Categorial factors have similar variance attribution to continuous proxies, but categorical factors attribute more variance to cell count and cell-type number.

The continuous substitution preserves the unconfounded structure of the L9 design and that it is observed that this pathway runs: DE difficulty → kNN purity → κ.

*Supplementary Figure 5 — Type III variance decomposition of Cohen's κ*

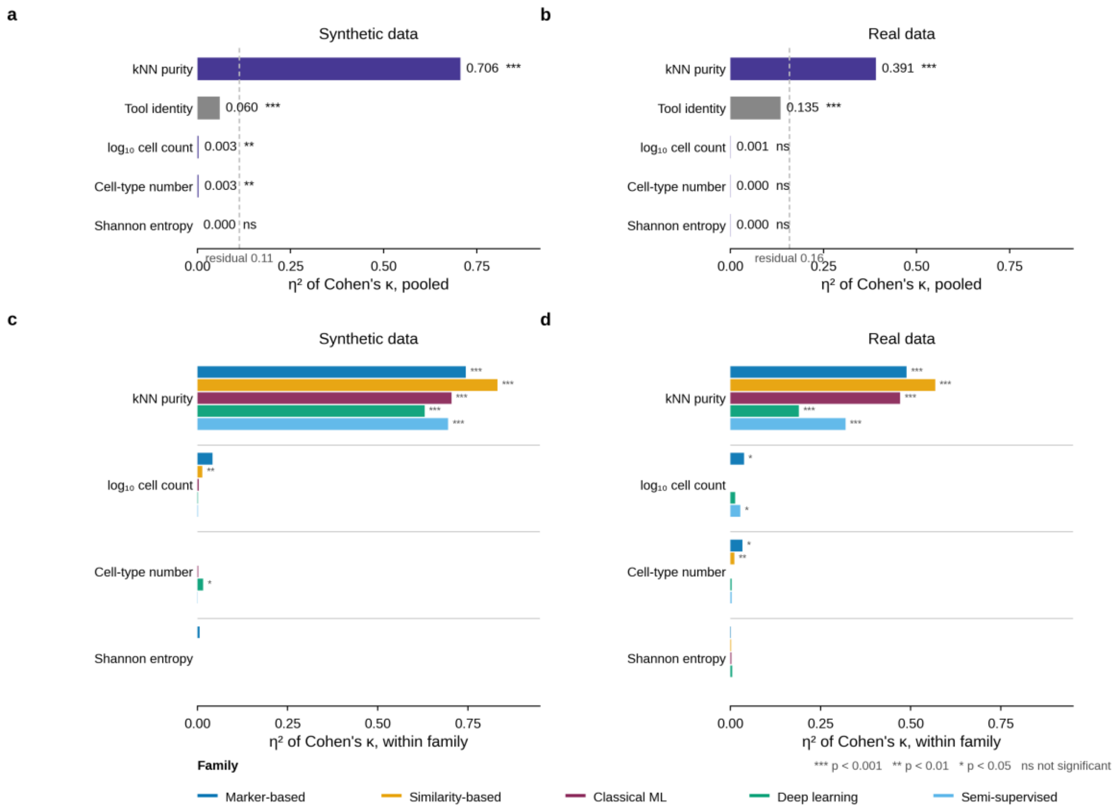


Type III η² of κ over the four continuous dataset variables ($\log_{10}$ cell count, cell-type number, Shannon entropy, kNN purity). **(a, b)** Pooled across all tools, with a tool-identity term. **(c, d)** The same decomposition fitted separately within each of the five cell-level families. Left column, synthetic data; right column, real data. The dashed line in (a) and (b) marks the residual.

Pooled, kNN purity carries the largest marginal $\eta^2$ on both data types (0.71 and 0.39) and tool identity a small share (0.06 and 0.14); each of the other three descriptors contributes little once purity is partialled out. Fitted within family, per-family purity $\eta^2$ spans 0.63–0.83 on the synthetic data and 0.19–0.57 on the real data, while the other three descriptors each contribute $\eta^2 \leq 0.05$ in every family on both data types.

| Family | Purity $\eta^2$, synthetic | Purity $\eta^2$, real |
|---|---|---|
| Similarity-based | 0.832 | 0.568 |
| Marker-based | 0.744 | 0.489 |
| Classical machine learning | 0.705 | 0.471 |
| Semi-supervised | 0.695 | 0.319 |

| Family | Purity $\eta^2$, synthetic | Purity $\eta^2$, real |
|---|---|---|
| Deep learning | 0.630 | 0.190 |

Purity’s dominance is not driven by any one family. Its attenuation from synthetic to real data is perhaps consistent with the additional biological noise present in real data. Purity has the greatest effect on similarity-based tools.

***Supplementary Figure 6 — Cluster-level reference: CIPR and cluster-level scmap***

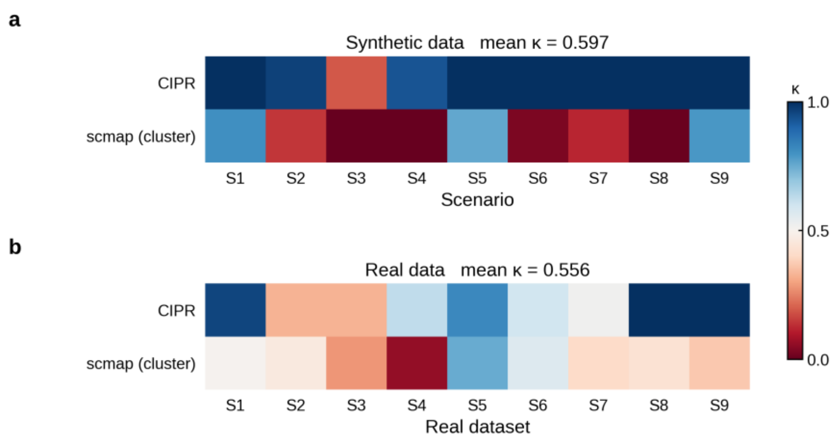


Per-tool mean κ for the two cluster-dependent similarity-based tools, CIPR and the cluster-level configuration of scmap, each supplied with a labeled reference and ground-truth cluster labels. **(a)** Nine synthetic scenarios, mean over three replicates. **(b)** Nine real datasets.

CIPR reaches a mean κ of 0.899 on the synthetic data scenarios and 0.686 on the real, against 0.295 and 0.425 for cluster-level scmap.

This records the accuracy achievable when CIPR and scmap operate at their native cluster-level granularity with perfect upstream clustering.

*Supplementary Figure 7 — Full marker family under derived markers*

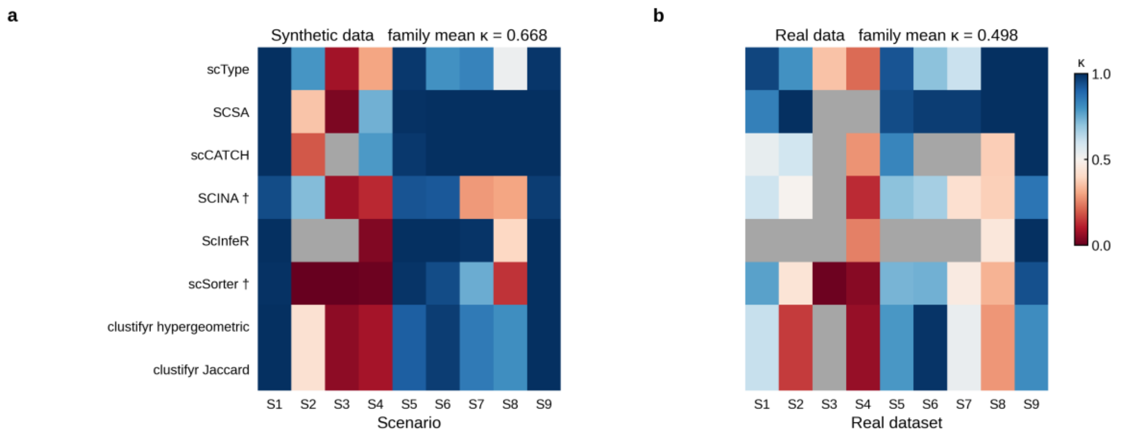


Per-tool κ heatmap for all eight marker-based tools (scType, SCSA, scCATCH, SCINA, ScInfeR, scSorter, clustifyr hypergeometric, clustifyr Jaccard) evaluated with derived marker lists (the top-20 differentially expressed genes per cell type from FindAllMarkers() on labeled data) and cluster labels, for the tools that require them. **(a)** Synthetic data: nine L9 scenarios, mean κ across three replicates. **(b)** Real data: nine within-platform datasets, one trial each. Gray indicates a failed run — 3 of 72 cells on the synthetic data and 14 of 72 on the real data.

Six of the eight tools are excluded from the cell-level comparison as they require cluster labels through unsupervised clustering which may confound the experimental design. Max performance is achieved here with idealized ground truth markers and clusters. SCINA and scSorter (flagged †), which do appear in the cell-level comparison and do not require clustering, are included here for completeness.

(a) Synthetic data, shows near-maximum performance on the four structurally easy scenarios (S1 0.993, S5 0.963, S6 0.953, S9 0.995) and a sharp decline on the hard-DE scenarios (S3 0.036, S4 0.271), consistent with the principal findings. SCSA (0.791), scCATCH (0.773) and scType (0.701) lead. The family mean across all eight tools and nine scenarios is κ = 0.668.

(b) Real data, performance shifts downward as the family mean drops to κ = 0.498, mostly due to the increased prevalence of failed runs on the real data, concentrated in ScInfeR. Over the cells that successfully produced annotations, the two means differ by only 0.08. SCSA (0.749) and scType (0.731) lead the real data ranking.

Further degradation observed under CellMarker 2.0 database marker-based annotation (Fig. 3e; family mean ontology score 0.438), can be interpreted as continued marker quality and mapping decline compared to the circumstances here.

***Supplementary Figure 8 — Cross-platform annotation transfer***

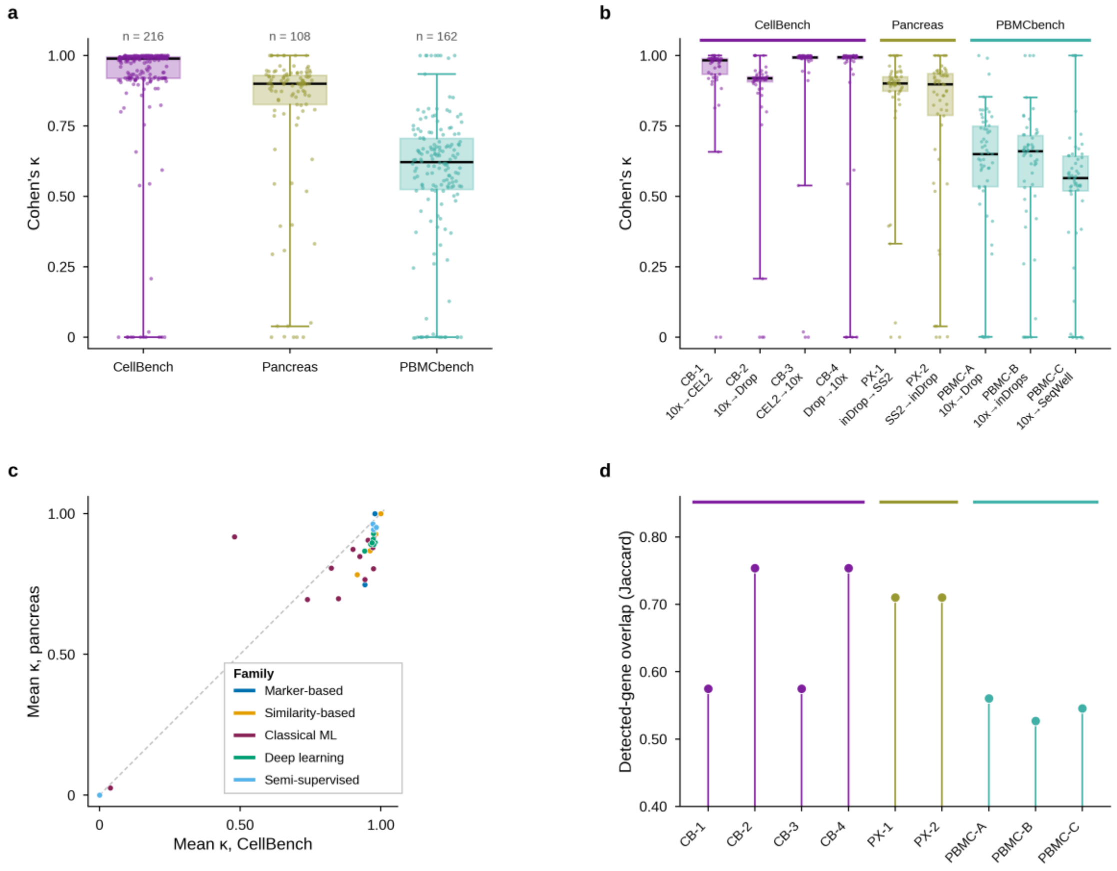


Cross-platform experiment, in which the reference and query come from different sequencing platforms but consist of a similar biological makeup. 55 tool configurations across nine trials in three blocks. **(a)** Cohen's κ by block. **(b)** Cohen's κ by trial, grouped by block; each trial is labeled with its reference → query platform pair. **(c)** Mean κ on the CellBench block against mean κ on the pancreas block. **(d)** Detected-gene overlap (Jaccard) between reference and query for each trial.

Median κ was 0.99 for CellBench ($n$ = 220), 0.90 for pancreas ($n$ = 110) and 0.62 for PBMCbench ($n$ = 165). Panels (a) and (b) show that the difference between blocks exceeds any within-block platform difference. Panel (c) shows that almost every tool sits near the diagonal, so the blocks separate tools together rather than reordering them significantly. Accuracy tracks the difficulty of the biological system rather than the platform contrast or the tool, which is the same pattern the within-platform experiments show. Fitting mean κ per trial against the query platform's kNN purity (Supplementary Table 2) gives $R^2 = 0.92$ and Spearman $\rho = 0.87$ ($p$ = 0.002, $n$ = 9 trials), so the separability–accuracy relationship holds in cross-platform annotation as well.

*Supplementary Figure 9 — Accuracy is uncorrelated with runtime or memory*

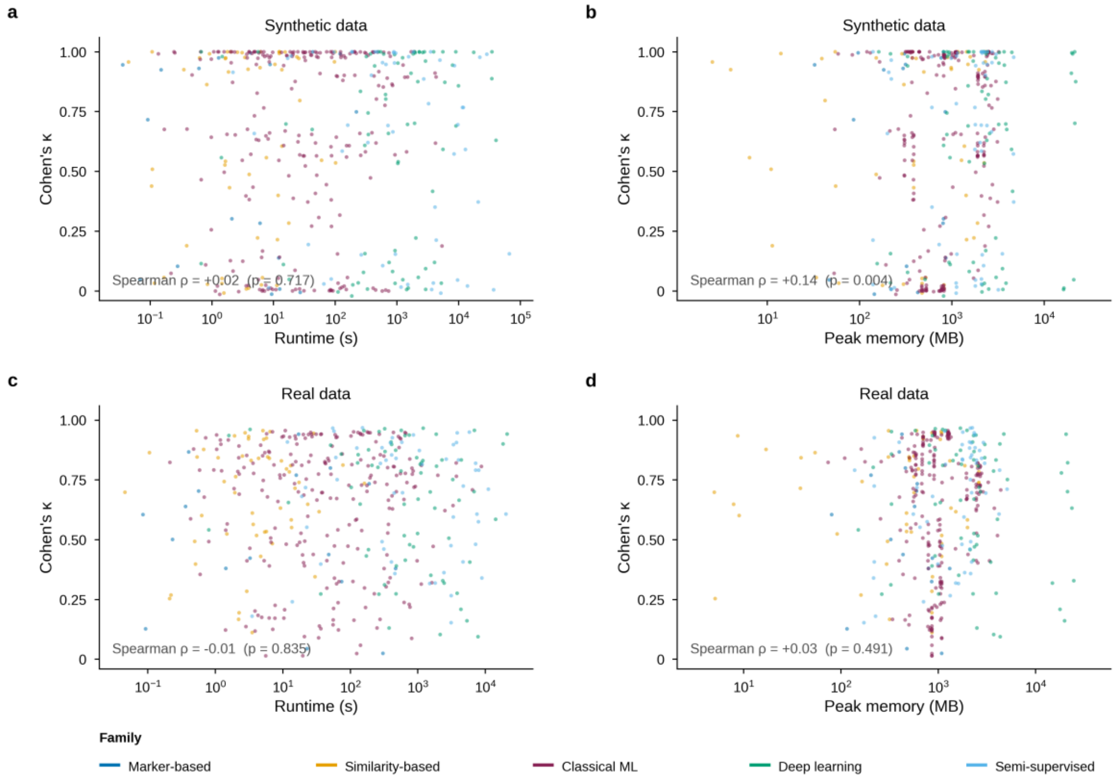


Scatter plots of per-(tool, dataset) mean κ against **(a)** $\log_{10}$ runtime and **(b)** $\log_{10}$ peak memory on the synthetic data, and **(c, d)** the corresponding real-data panels. Points are colored by family. Mean κ shows no meaningful association with either cost axis on either data type. Within the ranges tested, model complexity and resource cost do not predict annotation accuracy, and computationally cheap methods (such as similarity-based or LLM methods) matched or exceeded expensive families on the real data.

*Supplementary Figure 10 — Runtime and peak memory by family*

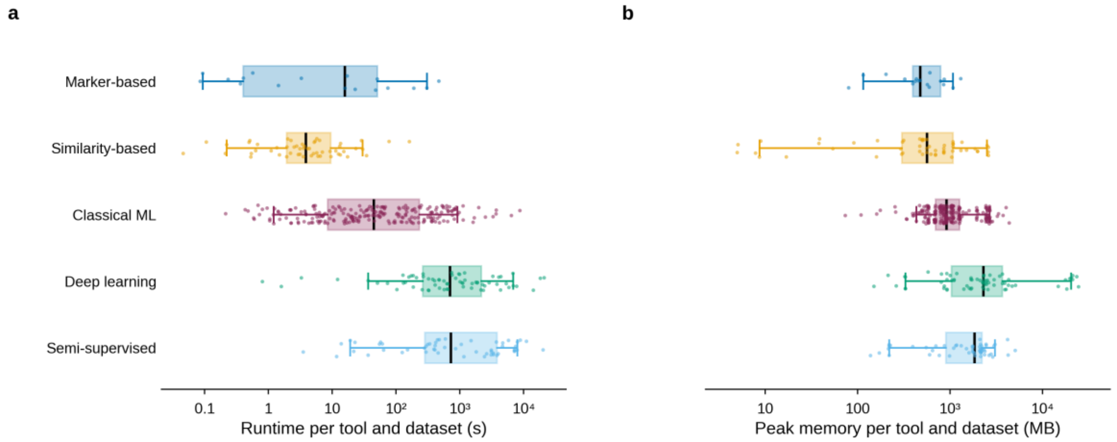


End to end runtime and peak memory per tool and dataset by method family on real data (n = 418). **(a)** Runtime. **(b)** Peak memory.

Families separate by orders of magnitude: marker-based and similarity-based methods complete in seconds to minutes, classical machine learning in minutes, and deep-learning and semi-supervised methods in tens of minutes to hours, with peak memory following the same ordering. Considering that the separability–accuracy relationship between families is near identical and linear, cost differentiates families based on accessibility and not on accuracy.

#### *Supplementary Figure 11 — Runtime and memory scaling with dataset size*

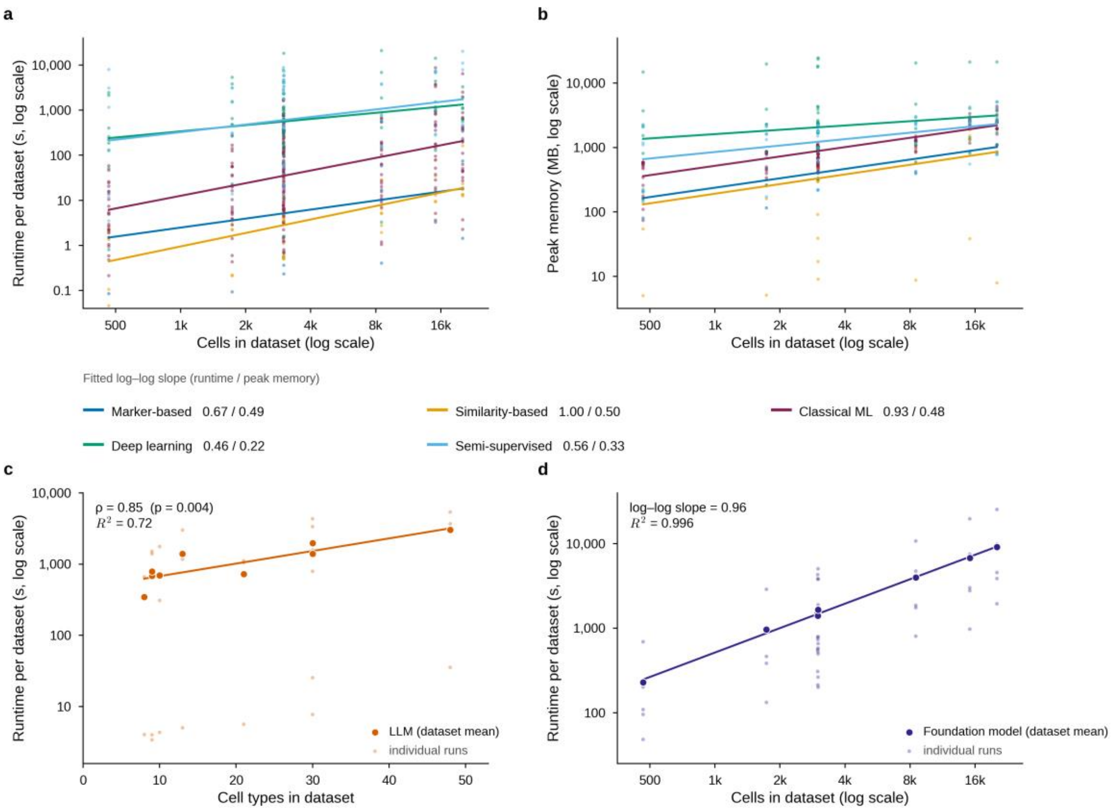


**(a)** $\log_{10}$ runtime against cell count, by family, fitted within family. **(b)** Same for $\log_{10}$ peak memory. Fitted log–log slopes, runtime / memory: similarity-based 1.00 / 0.50, classical ML 0.93 / 0.48, marker-based 0.67 / 0.49, semi-supervised 0.56 / 0.33, deep learning 0.46 / 0.22. **(c)** LLM runtime against number of cell types. **(d)** Foundation-model runtime against cell count. Faint points are individual runs, filled points are dataset means, and the fit utilizes the means.

For the LLM family, runtime scales with number of cell-types ($\rho = 0.85$, $p = 0.004$, $R^2 = 0.72$), which reflects the increasing inference calls needed for additional clusters. Foundation-model runtime scales near-linearly with cell count (log–log slope ≈ 0.96, $R^2 = 1.00$), spanning seconds to hours depending on model and dataset. Foundation-model peak GPU memory, by contrast, is set by the architecture rather than by dataset size and is mostly flat across datasets (not plotted).

#### *Supplementary Figure 12 — Per-tool runtime for the three derivative regimes*

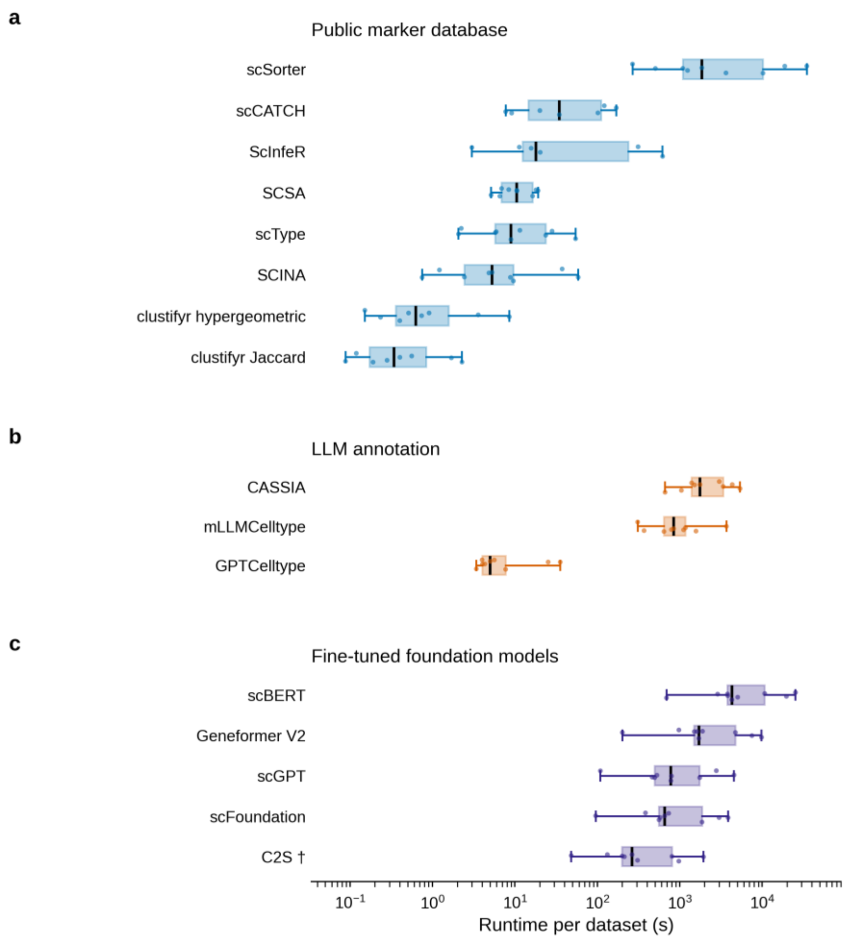


Per-tool runtime across the nine real datasets for **(a)** the eight database-driven marker tools, **(b)** the three LLM annotators and **(c)** the five fine-tuned foundation models. Boxes span the interquartile range, whiskers the full range, and a point is one dataset.

In (a), scSorter dominates at a median of 1,855 s while the remaining seven tools completed in a median of 0.3 to 35 s; the family's $\log_{10}$ cell-count dependence is driven almost entirely by scSorter, and the cluster-level tools run near instantaneously regardless of dataset size. In (b), the single-call GPTCelltype completed in 3.4–35.4 s per dataset while the multi-model consensus configurations mLLMCelltype and CASSIA required 309–3,685 s and 663–5,390 s respectively. LLM runtime is dependent on cell-type number and compounds in inference cost when querying several agents per cluster. In (c), runtime is combined train-plus-inference time; scBERT and Geneformer V2 are consistently the slowest per dataset on the L4 tier, and C2S and scFoundation complete fastest in their respective hardware contexts (Supplementary Method 3).

*Supplementary Figure 13 — Type III variance decomposition of log₁₀(runtime)*

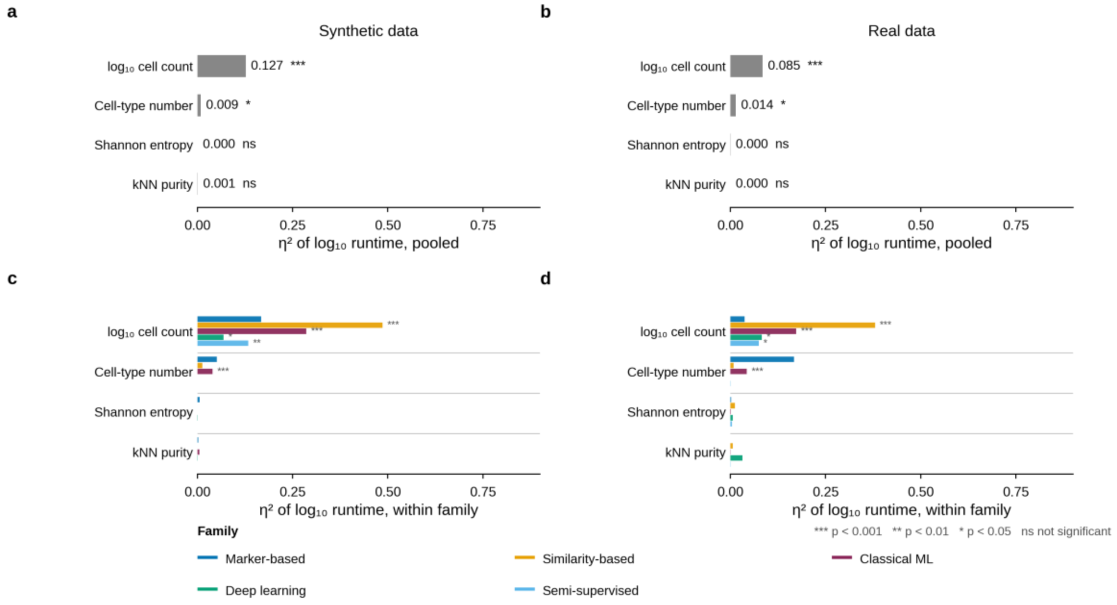


Type III $\eta^2$ for $\log_{10}$(runtime) across the four continuous variables. It is pooled and stratified by family on both data regimes. Cell count is the dominant driver of runtime for the similarity-based and classical machine-learning families but it is a much weaker contributor to runtime for the deep-learning and semi-supervised families. This is because their runtimes are dominated by training time rather than inference time.

The panel identifies which dataset properties drive computational cost within each family and confirms that the cell-count scaling reported in Supplementary Fig. 11 is carried principally by the similarity-based and classical machine-learning families.

### Supplementary Tables

*Supplementary Table 1 — Real-data panel: sources, protocol and label origin*

Sources for the nine real datasets.

| ID | Dataset | Species / tissue | Library type | Label origin | Source |
|---|---|---|---|---|---|
| S1 | Darmanis Brain 2015 [79] | Human cortex | Read-count (plate) | FACS isolation | Bioconductor scRNAseq |
| S2 | Marques Brain 2016 [80] | Mouse oligodendrocyte lineage | UMI | Author cluster annotation | Bioconductor scRNAseq |
| S3 | Nowakowski Cortex 2017 [46] | Human developing cortex | Read-count (plate) | Author cluster annotation | Original publication [46] |
| S4 | Grün Pancreas 2016 [81] | Human pancreatic islets | UMI | Author cluster annotation | Bioconductor scRNAseq |
| S5 | Tabula Muris FACS 3k 2018 [74] | Mouse multi-tissue | Read-count (plate) | FACS isolation | Bioconductor scRNAseq |
| S6 | He Skin 2020 [77] | Human skin | UMI | Author cluster annotation | Original publication [77] |
| S7 | Zhao Immune Fine 2020 [82] | Human immune subtypes | UMI | Author cluster annotation | Original publication [82]; Ensembl IDs mapped to HGNC symbols (Supplementary Method 1) |
| S8 | ZhengSort subset 2017 [3] | Human PBMC | UMI | Bead-based enrichment / FACS | Original publication [3] |

| ID | Dataset | Species / tissue | Library type | Label origin | Source |
|---|---|---|---|---|---|
| S9 | MacParland Liver Broad 2018 [45] | Human liver | UMI | Author cluster annotation | Bioconductor scRNAseq |

The cross-platform blocks draw on additional datasets sourced from their studies: CellBench [75] (10x v2 / CEL-seq2 / Drop-seq lung cancer cell-line mixtures), Baron [83] (inDrop) and Segerstolpe [84] (Smart-seq2) pancreatic islets, and PBMCbench [76] (10x v2 reference versus Drop-seq, inDrops and Seq-Well).

***Supplementary Table 2 — Cross-platform per-trial profile***

Dataset profiling for the cross-platform trial data.

| Trial | Block | Platform | Median lib. | Dropout (%) | Median genes | kNN purity | Mean DE strength π |
|---|---|---|---|---|---|---|---|
| CB-1 train | CellBench | 10x v2 | 109,992 | 45.02 | 9,073 | 0.997 | 431.4 |
| CB-1 test | CellBench | CEL-seq2 | 34,839 | 74.70 | 7,402 | 0.986 | 75.1 |
| CB-2 train | CellBench | 10x v2 | 109,992 | 45.02 | 9,073 | 0.997 | 431.4 |
| CB-2 test | CellBench | Drop-seq | 22,597 | 62.07 | 5,540 | 0.963 | 62.7 |
| CB-3 train | CellBench | CEL-seq2 | 34,839 | 74.70 | 7,402 | 0.986 | 75.1 |
| CB-3 test | CellBench | 10x v2 | 109,992 | 45.02 | 9,073 | 0.997 | 431.4 |
| CB-4 train | CellBench | Drop-seq | 22,597 | 62.07 | 5,540 | 0.963 | 62.7 |
| CB-4 test | CellBench | 10x v2 | 109,992 | 45.02 | 9,073 | 0.997 | 431.4 |

| Trial | Block | Platform | Median lib. | Dropout (%) | Median genes | kNN purity | Mean DE strength π |
|---|---|---|---|---|---|---|---|
| PX-1 train | Pancreas | Baron (inDrop) | 5,103 | 90.65 | 1,847 | 0.982 | 576.6 |
| PX-1 test | Pancreas | Segerstolpe (SS2) | 329,775 | 75.36 | 6,551 | 0.988 | 296.8 |
| PX-2 train | Pancreas | Segerstolpe (SS2) | 329,775 | 75.36 | 6,551 | 0.988 | 296.8 |
| PX-2 test | Pancreas | Baron (inDrop) | 5,103 | 90.65 | 1,847 | 0.982 | 576.6 |
| PBMC-A train | PBMCbench | 10x v2 (8 cl) | 2,708 | 97.51 | 1,112 | 0.927 | 217.6 |
| PBMC-A test | PBMCbench | Drop-seq (8 cl) | 1,436 | 97.15 | 756 | 0.854 | 146.5 |
| PBMC-B train | PBMCbench | 10x v2 (8 cl) | 2,708 | 97.51 | 1,112 | 0.927 | 217.6 |
| PBMC-B test | PBMCbench | inDrops (8 cl) | 806 | 97.98 | 469 | 0.834 | 166.8 |
| PBMC-C train | PBMCbench | 10x v2 (6 cl) | 2,701 | 97.53 | 1,109 | 0.949 | 265.3 |
| PBMC-C test | PBMCbench | Seq-well (6 cl) | 870 | 97.62 | 528 | 0.831 | 87.8 |

| Trial | Block | Reference | Query | Platform shift |
|---|---|---|---|---|
| CB-1 | CellBench | 10x v2 | CEL-seq2 | droplet → plate |

| Trial | Block | Reference | Query | Platform shift |
|---|---|---|---|---|
| CB-2 | CellBench | 10x v2 | Drop-seq | droplet → droplet |
| CB-3 | CellBench | CEL-seq2 | 10x v2 | plate → droplet |
| CB-4 | CellBench | Drop-seq | 10x v2 | droplet → droplet |
| PX-1 | Pancreas | Baron (inDrop) | Segerstolpe (Smart-seq2) | droplet → plate |
| PX-2 | Pancreas | Segerstolpe (Smart-seq2) | Baron (inDrop) | plate → droplet |
| PBMC-A | PBMCbench | 10x v2 (8 types) | Drop-seq | droplet → droplet |
| PBMC-B | PBMCbench | 10x v2 (8 types) | inDrops | droplet → droplet |
| PBMC-C | PBMCbench | 10x v2 (6 types) | Seq-Well | droplet → nanowell |

*Supplementary Table 3 — Master tool roster*

**Inputs supplied.** Each tool received the most informative version of the inputs it consumes, so that performance differences reflect algorithmic capability rather than input quality. The column entries denote:

- **Ref** — a labeled reference: a stratified training partition drawn from the same object as the test cells, sharing gene space, cell-type vocabulary, batch and class proportions.
- **Mkr** — derived markers: the top-20 differentially expressed genes per cell type from FindAllMarkers() on labeled data.
- **Clu** — supplied cluster labels: the ground-truth cell-type label vector passed as the tool's cluster argument, in place of an upstream unsupervised clustering step.
- **DB** — CellMarker 2.0, tissue-subsetted (Supplementary Method 2), supplied through each tool's native custom-marker interface.
- **FT** — a labeled training partition used for fine-tuning.
- **Ref†** — the same labeled reference, drawn from a different sequencing platform.
- **—** — the tool was not run in that setting.

| Tool | Family | Mechanism | Ref | Inputs | X-plat | DB / LLM | FM | Notes |
|---|---|---|---|---|---|---|---|---|
| SCINA | Marker | Semi-supervised EM assuming bimodal marker-gene expression | [13] | Mkr | Mkr | DB | — | Cell-level; rm_overlap=FALSE under the database (default TRUE strips genes from overlapping CellMarker 2.0 signatures) |
| scSorter | Marker | Marker-guided joint model of marker genes + HVGs | [47] | Mkr | Mkr | DB | — | Cell-level |
| scType | Marker | Cell-type specificity scoring (positive + negative markers) | [12] | Mkr + Clu | Mkr + Clu | DB + Clu | — | Cluster-dependent; excluded from the cell-level comparison |
| scCATCH | Marker | Evidence-based cluster scoring vs tissue-specific database | [48] | Mkr + Clu | Mkr + Clu | DB + Clu | — | Cluster-dependent; excluded from the cell-level comparison |
| SCSA | Marker | DEG identification + confidence-weighted database scoring | [49] | Mkr + Clu | Mkr + Clu | DB + Clu | — | Cluster-dependent; also queries bundled database alongside CellMarker 2.0 under the database; excluded from the cell-level comparison |

| Tool | Family | Mechanism | Ref | Inputs | X-plat | DB / LLM | FM | Notes |
|---|---|---|---|---|---|---|---|---|
| clustifyr (hypergeometric) | Marker | Cluster enrichment against reference marker sets | [50] | Mkr + Clu | Mkr + Clu | DB + Clu | — | Cluster-dependent; excluded from the cell-level comparison |
| clustifyr (Jaccard) | Marker | Cluster Jaccard similarity against reference marker sets | [50] | Mkr + Clu | Mkr + Clu | DB + Clu | — | Cluster-dependent; excluded from the cell-level comparison |
| ScInfeR | Marker | Marker annotation refined by kNN-weighting in cell neighborhood graph (requires UMAP of test data) | [51] | Mkr + Clu | Ref † + Clu | DB | — | Cluster-dependent under derived markers; excluded from the cell-level comparison |
| CIPR | Similarity | Dot-product comparison of DE log-fold-changes between query clusters and reference | [55] | Ref + Clu | Ref † + Clu | — | — | Cluster-dependent; reported in Supplementary Fig. 6 only |
| CHETAH | Similarity | Hierarchical classification tree traversed by correlation score | [56] | Ref | Ref † | — | — | Included |
| SingleR | Similarity | Spearman correlation to reference profiles with marker-based fine-tuning | [14] | Ref | Ref † | — | — | Included |
| Seurat label transfer (PCA) | Similarity | PCA-anchor integration with anchor-weighted label projection | [52, 53] | Ref | Ref † | — | — | Included |

| Tool | Family | Mechanism | Ref | Inputs | X-plat | DB / LLM | FM | Notes |
|---|---|---|---|---|---|---|---|---|
| Seurat label transfer (RPCA) | Similarity | RPCA-anchor integration with anchor-weighted label projection | [52, 53] | Ref | Ref † | — | — | Included |
| Seurat label transfer (CCA) | Similarity | CCA-anchor integration with anchor-weighted label projection | [52, 53] | Ref | Ref † | — | — | Included |
| scmap (cell) | Similarity | Cell-level nearest-neighbor projection in cosine-distance space | [15] | Ref | Ref † | — | — | Included (cell-level configuration) |
| scmap (cluster) | Similarity | Cluster-level nearest-neighbor projection in cosine-distance space | [15] | Ref + Clu | Ref † | — | — | Cluster-dependent; reported in Supplementary Fig. 6 only |
| scibetR | Classical ML | Multinomial model with maximum-entropy gene selection | [54] | Ref | Ref † | — | — | Included |

| Tool | Family | Mechanism | Ref | Inputs | X-plat | DB / LLM | FM | Notes |
|---|---|---|---|---|---|---|---|---|
| scPred (×17 configs) | Classical ML | Feature-projection classifier: default radial-kernel SVM + 16 caret algorithms (avNNet, xgbTree, random forest, GLM, glmboost, LDA, kNN, linear SVM, polynomial SVM, naïve Bayes, Bayesian GLM, MARS-style earth, MLP, neural network, regularized logistic, multinomial) | [16] | Ref | Ref † | — | — | Included; AdaBoost omitted (κ ≈ 0 across scenarios) |
| scClassify | Classical ML | Ensemble of weighted kNN with hierarchical cell-type trees | [57] | Ref | Ref † | — | — | Included |
| singleCellNet | Classical ML | Random forest with top-scoring-pair feature transformation | [17] | Ref | Ref † | — | — | Included |
| CellTypist | Classical ML | Multinomial logistic regression with SGD training | [18] | Ref | Ref † | — | — | Included; Python |
| scAnnotate | Classical ML | Ensemble of zero-inflated mixture models | [20] | Ref | Ref † | — | — | Included; Python |
| CaSTLe | Classical ML | XGBoost on discretized expression with mutual-information-based feature selection | [58] | Ref | Ref † | — | — | Included |

| Tool | Family | Mechanism | Ref | Inputs | X-plat | DB / LLM | FM | Notes |
|---|---|---|---|---|---|---|---|---|
| scID | Classical ML | Fisher's linear discriminant analysis with cluster-specific signatures | [19] | Ref | Ref † | — | — | Included |
| scLearn | Classical ML | Discriminative component analysis (DCA) with M3Drop feature selection | [59] | Ref | Ref † | — | — | Included |
| NeuCA | Deep learning | Hierarchical feedforward neural network | [60] | Ref | Ref † | — | — | Included |
| ACTINN | Deep learning | Multilayer perceptron | [21] | Ref | Ref † | — | — | Included |
| Cell BLAST | Deep learning | Biologically-informed variational autoencoder with adversarial regularization | [61] | Ref | Ref † | — | — | Included |
| CIForm | Deep learning | Transformer-based classifier | [62] | Ref | Ref † | — | — | Included |
| scBalance | Deep learning | Sparse neural network with adaptive weight sampling for rare types | [63] | Ref | Ref † | — | — | Included |
| scDeepHash | Deep learning | Deep hashing model for cell retrieval | [64] | Ref | Ref † | — | — | Included |
| scMMT | Deep learning | Multi-modal transformer applied in RNA-only mode | [65] | Ref | Ref † | — | — | Included |

| Tool | Family | Mechanism | Ref | Inputs | X-plat | DB / LLM | FM | Notes |
|---|---|---|---|---|---|---|---|---|
| TOSICA | Deep learning | Multi-head self-attention transformer with pathway-attention interpretability | [22] | Ref | Ref † | — | — | Included |
| CAMLU | Semi-supervised | Autoencoder reconstruction error with iterative bimodal feature selection, paired with an SVM for known cell types | [66] | Ref | Ref † | — | — | Included; consumes unlabeled test data |
| MARS | Semi-supervised | Meta-learning of a shared embedding with cell-type landmarks | [67] | Ref | Ref † | — | — | Included; consumes unlabeled test data |
| scANVI | Semi-supervised | Semi-supervised deep generative model extending scVI | [23] | Ref | Ref † | — | — | Included; consumes unlabeled test data |
| scArches | Semi-supervised | Architecture-surgery fine-tuning of pre-trained scVI/scANVI models | [24] | Ref | Ref † | — | — | Included; consumes unlabeled test data |
| scSemiCluster | Semi-supervised | Semi-supervised domain adaptation with structural regularization | [68] | Ref | Ref † | — | — | Included; consumes unlabeled test data |
| scSemiGAN | Semi-supervised | Semi-supervised generative adversarial annotation | [69] | Ref | Ref † | — | — | Included; consumes unlabeled test data |

| Tool | Family | Mechanism | Ref | Inputs | X-plat | DB / LLM | FM | Notes |
|---|---|---|---|---|---|---|---|---|
| scBERT | FM | Bidirectional encoder-only transformer (~1M cells; Performer attention) | [25] | — | — | — | FT | Foundation-model experiment only; fine-tuning; NVIDIA L4 |
| scGPT | FM | Generative decoder-style transformer (~33M cells; FlashAttention) | [27] | — | — | — | FT | Foundation-model experiment only; fine-tuning; NVIDIA L4 |
| Geneformer V2 | FM | Rank-tokenization transformer (~104M transcriptomes) | [26] | — | — | — | FT | Foundation-model experiment only; fine-tuning; NVIDIA L4 |
| scFoundation | FM | Linear-complexity-attention transformer (~50M cells) | [28] | — | — | — | FT | Foundation-model experiment only; fine-tuning; NVIDIA L4 |
| C2S (Cell2Sentence) | FM | Autoregressive language model on cell-as-sentence encoding (Pythia-410M backbone) | [29] | — | — | — | FT | Foundation-model experiment only; fine-tuning; NVIDIA RTX PRO 6000 (102 GB VRAM) |
| GPTCelltype | LLM | Single-model prompting with per-cluster top DE markers (GPT-5.4) | [30] | — | — | Clu | — | Database and LLM experiments only |

| Tool | Family | Mechanism | Ref | Inputs | X-plat | DB / LLM | FM | Notes |
|---|---|---|---|---|---|---|---|---|
| CASSIA | LLM | Multi-agent pipeline with iterative refinement (GPT-5.4 primary; Claude Sonnet 4.6 scoring/annotation-boost) | [31] | — | — | Clu | — | Database and LLM experiments only |
| mLLMCelltype | LLM | Multi-model consensus (GPT-5.4 + Claude Sonnet 4.6 + Gemini 3 Pro) with uncertainty quantification | [32] | — | — | Clu | — | Database and LLM experiments only |

- 
- 

scPred contributes 17 configurations: the default radial-kernel SVM plus 16 additional caret models (avNNet, xgbTree, random forest, GLM, glmboost, LDA, kNN, linear SVM, polynomial SVM, naïve Bayes, Bayesian GLM, MARS-style earth, MLP, neural network, regularized logistic, multinomial).

### Supplementary Methods

#### *Supplementary Method 1 — Gene identifier alignment*

For real datasets, gene names were formatted into standard human gene-symbol convention.

The Zhao Immune Fine dataset (S7) was distributed with Ensembl gene identifiers and these were converted to HGNC-approved symbols via the EnsDb.Hsapiens.GRCh38 annotation retrieved through AnnotationHub. Unmapped Ensembl IDs were dropped and duplicate symbols were collapsed. The remaining seven human datasets (Darmanis S1, Nowakowski S3, Grün S4, He-Skin S6, ZhengSort S8, MacParland S9, and the human cell-line and PBMC blocks used in the cross-platform experiment) were distributed with HGNC symbols and did not require mapping.

The two mouse datasets — the Marques oligodendrocyte panel (S2) and the Tabula Muris FACS panel (S5) — were case-folded to uppercase mouse symbols (e.g. *Pdgfra* → *PDGFRA*) and rodent-specific gene entries carrying the Rik suffix were discarded. This case-folding is an approximate alignment to the human symbol convention rather than a formal one-to-one ortholog mapping. The case-folded mouse vocabularies recovered a per-dataset gene-overlap fraction similar to the fraction of foundation-model vocabulary recovered compared to the human-source datasets. This affects database-connected marker tools, LLM pipelines, and foundation models. Cell-level supervised tools were applied to the top 2,000 highly variable genes selected within each dataset and are unaffected.

***Supplementary Method 2 — CellMarker 2.0 tissue subsetting***

For each of the nine real datasets, CellMarker 2.0 was subsetted to the tissue(s) of the input dataset before scoring as to avoid inflated scoring against a full marker database.

| Dataset ID | Dataset | Tissue subset applied |
|---|---|---|
| S1 | Darmanis Brain 2015 | Brain |
| S2 | Marques Brain 2016 | Brain |
| S3 | Nowakowski Cortex 2017 | Brain + Fetal brain |
| S4 | Grün Pancreas 2016 | Pancreas |
| S5 | Tabula Muris FACS 3k 2018 | Blood, Bone Marrow, Brain, Skin, Epidermis, Bladder, Bronchus, Thymus, Heart, Colon, Adipose Tissue, Skeletal Muscle, Pancreas |
| S6 | He Skin 2020 | Skin |
| S7 | Zhao Immune Fine 2020 | Blood |
| S8 | ZhengSort 20k 2017 | Blood |
| S9 | MacParland Liver Broad 2018 | Liver |

The subset was supplied to each tool through its native custom-marker parameter.

#### *Supplementary Method 3 — C2S (Cell2Sentence) architecture and hardware*

The C2S architecture is a hybrid of both foundation-model and LLM families. It applies a generative language-model backbone (Pythia-410M) to expression data rendered as cell sentences in contrast to the embedding-based backbones (scBERT, scGPT, scFoundation, Geneformer V2).

Additionally, the four embedding-based foundation models were fine-tuned from task-agnostic self-supervised weights, while C2S was fine-tuned from an annotation-task-specific checkpoint (C2S-Pythia-410m-cell-type-prediction). C2S required substantially more GPU memory and ran on the NVIDIA RTX PRO 6000 Blackwell Server Edition (102 GB VRAM) rather than the NVIDIA L4 (22.5 GB VRAM) used for the other four. Absolute runtime comparisons between C2S and the other foundation models should be read with this hardware difference in mind.

***Supplementary Method 4 — Orthogonality retention and the categorical decomposition***

The L9 array sets four properties at discrete levels but the real datasets were profiled not novelly designed. Substituting measured continuous quantities ($\log_{10}$ total cells, number of cell types, normalized Shannon entropy, mean kNN purity) for the discrete design variables puts risk that the proxies correlate with each other.

The substitution holds as across the nine synthetic scenarios, the largest pairwise correlation between proxies is $r = 0.36$. The correlation between cell count cell-type number, in particular, is $r = 0.00$. Variance inflation factors were calculated to be low at 1.15–1.19 ($\log_{10}$ cells 1.158, cell types 1.152, entropy 1.152, purity 1.194) (Supplementary Fig. 4a).

For the synthetic data we also decomposed κ across the four design factors at their actual categorical levels — cell count {500, 3,000, 15,000}, imbalance {balanced, mild, severe}, cell types {5, 15, 30}, DE difficulty {easy, medium, hard} and observed a similar variance attribution to the continuous proxies (Supplementary Fig 4c).

### Supplementary Notes

***Supplementary Note 1 — MCC concordance with Cohen's κ***

The multi-class Matthews correlation coefficient (MCC, Gorodkin formulation) can be utilized for imbalanced classification because it summarizes the full confusion matrix symmetrically. It proved volatile here because of the high unassigned rates some tools generate on harder datasets. Cohen's κ is robust to this issue and has historical use precedent [11].

Still, κ and MCC were highly concordant across experiments: the two tracked each other closely, with only a small number of outliers exceeding $|\kappa - \mathrm{MCC}| > 0.05$ and no systematic directional bias. Tool rankings and family-level conclusions are unchanged when MCC is substituted for κ.

*Supplementary Note 2 — Extended limitations and scope*

**Orthogonal main effects only.** The type III Taguchi L9 orthogonal design estimates the main effects independently, but cannot resolve nonlinear interactions among the four design factors. Capturing interactions would require a larger array. Our variance decomposition should be interpreted as a main-effects model within defined parameter ranges.

**Idealized synthetic geometry.** Splatter's probabilistic model treats genes as independently differentially expressed, failing to capture correlated gene-regulatory networks, multi-donor variance, or complex batch confounding. Synthetic datasets can consequently exhibit cleaner population separation than real biological datasets at equivalent fold changes (Supplementary Fig. 2) [37]. While this geometric idealization inflates absolute performance, general cross-tool comparisons remain valid and consistent through real data validation.

**Bounded cell-count window.** The cell-count range was chosen based on compute and memory constraints. Further testing is needed to establish whether these findings extend to larger cell-count ranges.

**L9 matching over provenance.** The real-data validation panel prioritizes matching to the L9 design over annotation provenance. Gold-standard isolation methods (e.g. FACS sorting or bead-based enrichment) are increasingly rare; most public atlases [4] carry computationally derived annotations which introduces circular validation.

**Limited foundation-model sample.** The foundation-model experiment was restricted to five specific pre-trained architectures. The broader structural question of whether self-supervised pre-training provides general and scalable transfer learning remains unanswered and is a current area of study.

**Foundation-model evaluation with fine-tuning only.** We evaluate the foundation models adapted by full fine-tuning; zero-shot performance is not formally benchmarked against our baselines. Informal testing was consistent with recent literature showing that untuned zero-shot cell-type annotation yields performance near or below chance baselines [33, 34].

**Matched reference conditions.** All reference-consuming tools in the cell-level comparison received a reference derived from a stratified split of the query dataset. This reference shared identical gene space, cell-type vocabularies, batch composition, and class proportions with the test partition. Because real-world deployment relies on independently curated atlases with inevitable batch effects, proportional skews, and vocabulary mismatches, these performance estimates represent theoretical upper limits. The cross-platform experiment and the database experiment are meant to test annotation tools with less input control and more deployment realism.

**Supplied cluster labels.** The cluster-consuming tools in the database and LLM experiments received the true cell-label vector (true clusters) as their input, rather than the output of an upstream unsupervised algorithm. In real pipelines, Louvain and Leiden clustering introduce confounding error. Performance metrics for these cluster-dependent tools are therefore upper bounds conditioned on perfect upstream clustering.

**Single-split uncertainty on the real data.** Unlike the three-seed replicates used in the synthetic data experiments, the real-data panel used a single stratified 80/20 split per dataset. Real accuracy scores are therefore point estimates that lack tool-specific standard errors, and cross-family comparisons rely on a single observation per tool–dataset pair.

**LLM ontology scoring without human-rater calibration.** The three-model language panel's scores have not been calibrated against a human-expert panel, so absolute values should be interpreted strictly as automated relative metrics rather than as human-concordant ones. Some scorers share a model family with the tools they score.

**Semi-supervised structural advantages.** The six semi-supervised tools have a structural advantage over reference-only classifiers: they access the unlabeled test data distribution during model fitting. This joint estimation allows them to adapt directly to the query distribution. This non-equivalent information regime explains part of the semi-supervised family's high real-data mean ($\kappa = 0.675$).

**Pre-training corpus data leakage risk.** The pre-training corpora for scGPT and scFoundation included portions of the CZ CELLxGENE repository [5], creating a potential risk of data leakage if any of our evaluation datasets were included in their training sets. Geneformer V2 pre-training may also have data overlap, whereas scBERT (trained on PanglaoDB [73]) carries no such risk.

***Supplementary Note 3 — Runtime measurement and the kNN-purity as a diagnostic deployment heuristic***

**Runtime measurement.** Because our benchmarking harness records model training and inference within a single execution call, the high absolute runtime of the classical machine-learning, deep-learning and semi-supervised families is dominated by per-dataset model fitting rather than by annotation inference. Similarity- and marker-based tools carry almost no training overhead. During deployment, once a supervised model has been trained on a reference atlas, inference on incoming query cells is substantially faster. This narrows the performance-to-cost gap between the trained-model families and the reference-free alternatives, and supports the use of pre-trained tissue libraries to bypass training overhead entirely, with the tradeoff of a rigid cell-type vocabulary.

**kNN-purity as a diagnostic.** Practitioners can consider using a ground truth reference set similar to the query set to obtain kNN purity and predict automatic annotation performance. Without a ground-truth reference, practitioners could use the kNN purity computed from unsupervised clustering on the query set. Empirically, the relationship between the two followed a pattern:

- **Structurally clear datasets.** Ground-truth purity sat slightly above the unsupervised-clustering value.
- **Moderately difficult datasets.** Ground-truth purity sat slightly below the unsupervised value, with minor variation in the absolute gap.
- **Severe stress-test datasets.** Ground-truth purity substantially exceeded the unsupervised-clustering value.

In our experience this last divergence signals poor dataset or annotation quality rather than any structure (i.e. Nowakowski Cortex and Grün Pancreas). An unsupervised-clustering set's purity should therefore be read as an approximate-to-slightly- conservative proxy on easy-to-moderate data. Manual annotation or refinement is needed in the extreme/hard dataset case.
